# AP-1 transcription factor network explains diverse patterns of cellular plasticity in melanoma cells

**DOI:** 10.1101/2021.12.06.471514

**Authors:** Natacha Comandante-Lou, Douglas G. Baumann, Mohammad Fallahi-Sichani

**Author notes:** These authors contributed equally (Natacha Comandante-Lou, Douglas G. Baumann).

## Abstract

Cellular plasticity associated with fluctuations in transcriptional programs allows individual cells in a tumor to adopt heterogeneous differentiation states and switch phenotype during their adaptive responses to therapies. Despite increasing knowledge of such transcriptional programs, the molecular basis of cellular plasticity remains poorly understood. Here, we combine multiplexed transcriptional and protein measurements at population and single-cell levels with multivariate statistical modeling to show that the state of AP-1 transcription factor network plays a unifying role in explaining diverse patterns of plasticity in melanoma. We find that a regulated balance between AP-1 factors cJUN, JUND, FRA2, FRA1 and cFOS determines the intrinsic diversity of differentiation states and adaptive responses to MAPK inhibitors in melanoma cells. Perturbing this balance through genetic depletion of specific AP-1 proteins, or by MAPK inhibitors, shifts cellular heterogeneity in a predictable fashion. Thus, AP-1 may serve as a critical node for manipulating cellular plasticity with potential therapeutic implications.

## Introduction

Individual cells, even those derived from the same clone, respond heterogeneously to environmental perturbations (Mitchell and Hoffmann, 2018; Munsky et al., 2012). Nongenetic heterogeneity can arise due to variances associated with transcriptional state plasticity (Battich et al., 2015; Gupta et al., 2011; Munsky et al., 2012; Symmons and Raj, 2016). Although such plasticity is required for the proper development of complex organisms (Arias and Hayward, 2006), it limits the efficacy of therapies that target abnormally-activated signaling pathways (Boumahdi and de Sauvage, 2020; Sharma et al., 2010). An example of cell-to-cell transcriptional heterogeneity with phenotypic consequences for therapy resistance is observed in melanomas (Emert et al., 2021; Fallahi Sichani et al., 2017; Shaffer et al., 2017). Numerous studies have associated fluctuations in the state of MAPK inhibitor sensitivity across BRAF-mutant melanoma cells to intrinsic variations in their differentiation state (Baron et al., 2020; Belote et al., 2021; Khaliq et al., 2021; Rambow et al., 2018; Tsoi et al., 2018; Wouters et al., 2020). The reported heterogeneity spans a range of transcriptionally distinguishable states, including a melanocytic phenotype that expresses melanocyte lineage markers SOX10 and MITF (Lin and Fisher, 2007), to less drug-sensitive states, including neural crest-like cells that express NGFR (Fallahi Sichani et al., 2017; Mica et al., 2013), and innately drug-resistant, undifferentiated cells characterized by the overexpression of AXL and loss of SOX10 and MITF (Konieczkowski et al., 2014; Müller et al., 2014). In addition to intrinsic disparities in differentiation state, drug-induced responses may help a fraction of cells rewire their state of MAPK inhibitor sensitivity, most commonly through adaptive changes in differentiation state (Fallahi Sichani et al., 2017; Marin-Bejar et al., 2021; Rambow et al., 2018; Smith et al., 2016) or via reactivation of the MAPK pathway (Gerosa et al., 2020; Lito et al., 2012). Although the emergence and consequences of such intrinsic and adaptive heterogeneities are widely recognized, there is still more to learn about their origins and possible connection at a molecular level. For example, it is unclear whether these seemingly distinct forms of heterogeneity arise from independent mechanisms, or whether the observed variability in the initial state of cells and adaptive changes following MAPK inhibitor treatment could both be traced back to a common subset of molecular players.

Transcription factor networks that regulate the expression of genes in response to signaling pathway perturbations play a key role in creating the biological noise that leads to population heterogeneity (Huang, 2009; Pedraza and van Oudenaarden, 2005). The AP-1 protein family comprises one such network that serves as a major transcription node, integrating inputs from the upstream MAPK signaling pathway (Karin, 1995). In addition to linking signal transduction to transcription, AP-1 proteins have been recently identified to serve as pioneer factors, establishing chromatin states that predispose cells to transcriptional programs driven by other transcription factors or histone modifications, thereby guiding cells towards paths of differentiation or epigenetic reprogramming (Madrigal and Alasoo, 2018; Martínez-Zamudio et al., 2020; Phanstiel et al., 2017; Vierbuchen et al., 2017). These roles are consistent with numerous reports on AP-1 proteins being involved in resistance to MAPK inhibitors, cell state heterogeneity, and therapy-induced dedifferentiation in melanomas and other cancers (Emmons et al., 2019; Fallahi-Sichani et al., 2015; Fallahi Sichani et al., 2017; Haas et al., 2021; Johannessen et al., 2013; Kong et al., 2017; Maurus et al., 2017; Ramsdale et al., 2015; Riesenberg et al., 2015; Torre et al., 2021; Wouters et al., 2020). Despite these reports, we lack a clear understanding of the rules that define AP-1 behavior and its role in explaining the intrinsic plasticity and the diversity of adaptive responses to MAPK signaling perturbations. This gap in our knowledge may be addressed by a system-wide analysis with single-cell precision to reveal interdependencies between an array of AP-1 proteins, which comprise over a dozen transcription factors, including JUN, FOS, and ATF subfamilies (Rodríguez-Martínez et al., 2017), their post-translational modification states, and their association with melanoma cell phenotypes at a single-cell level.

In this paper, we test the hypothesis that the state of the AP-1 transcription factor network determines the intrinsic diversity of phenotypic states (i.e., differentiation states) and drug-induced changes in MAPK signaling in BRAF-mutated melanoma cells. We define the AP-1 state as the combinatorial concentrations of AP-1 proteins, their phosphorylation state, and their transcriptional activity, which are either measurable experimentally, or inferable by using bioinformatics tools. Our systems biology approach combines multiplexed measurements of the AP-1 state, MAPK signaling activity and differentiation state, at population and single-cell levels, across many genetically characterized melanoma cell lines before and after their exposure to BRAF/MEK inhibitors. We apply statistical learning to capture the predictivity of AP-1 states, and corresponding AP-1 factors, for phenotypic heterogeneity in melanoma cultures and patient-derived tumors. We then employ RNAi-mediated knockdown experiments to validate the causality of our statistical predictions in heterogeneous melanoma cell populations. We find that a tightly regulated balance between AP-1 transcription factors cJUN, JUND, FRA2, FRA1 and cFOS and their transcriptional activity determines the baseline differentiation state of melanoma cells. This balance is perturbed following MAPK pathway inhibition. Nevertheless, MAPK-inhibitor-induced changes in the AP-1 state, including the abundance of cJUN and its phosphorylation, as well as the phosphorylation state of FRA1, remain strong predictors of drug-induced changes in differentiation state and the efficiency of MAPK pathway inhibition, respectively. These results show that the state of AP-1 network offers a critical transcriptional context, which controls not only the initial state of melanoma cells and their population heterogeneity, but also their adaptive changes immediately following MAPK pathway inhibition.

## Results

### Single-cell AP-1 protein levels predict differentiation state heterogeneity in melanoma cells

To quantify the baseline heterogeneities in differentiation state and to assess their covariation with AP-1 proteins across genetically diverse or isogenic melanoma cell populations, we utilized an iterative indirect immunofluorescence imaging (4i) protocol (Gut et al., 2018) in conjunction with high-throughput automated microscopy (Figure 1A). We multiplexed measurements of 21 proteins using 4i-validated antibodies in 19 BRAF-mutant melanoma cell lines (Figure 1B). The measurements included total levels of eleven AP-1 transcription factors (including cFOS, FRA1, FRA2, cJUN, JUNB, JUND, ATF2, ATF3, ATF4, ATF5 and ATF6), six AP-1 phosphorylation states (including p-cFOS^S32^, p-FRA1^S265^, p-cJUN^S73^, p-ATF1^S63^, p-ATF2^T71^ and p-ATF4^S245^), and four differentiation state markers MITF, SOX10, NGFR and AXL. Importantly, these four differentiation state markers were previously reported to represent transcriptionally distinct melanoma differentiation states (Khaliq et al., 2021; Tsoi et al., 2018). The panel of 19 melanoma cell lines tested represented a broad spectrum of differentiation states, including populations of melanocytic (MITF^High^ /SOX10^High^ /NGFR^Low^ /AXL^Low^), transitory (MITF^High^ /SOX10^High^ /NGFR^High^ /AXL^Low^), neural crest-like (MITF^Low^ /SOX10^High^ /NGFR^High^ /AXL^High^) and undifferentiated (MITF^Low^ /SOX10^Low^ /NGFR^Low^ /AXL^High^) cells (Figures 1C, S1A). We and others have shown that the frequency of these states in melanoma cell populations varies from one tumor to another and predicts their overall sensitivity to MAPK inhibitors (Khaliq et al., 2021; Rambow et al., 2018; Tirosh et al., 2016). Here, we asked whether the observed heterogeneities in differentiation state could be explained by variations in patterns of AP-1 measurements at a single-cell level.

**Figure 1.**
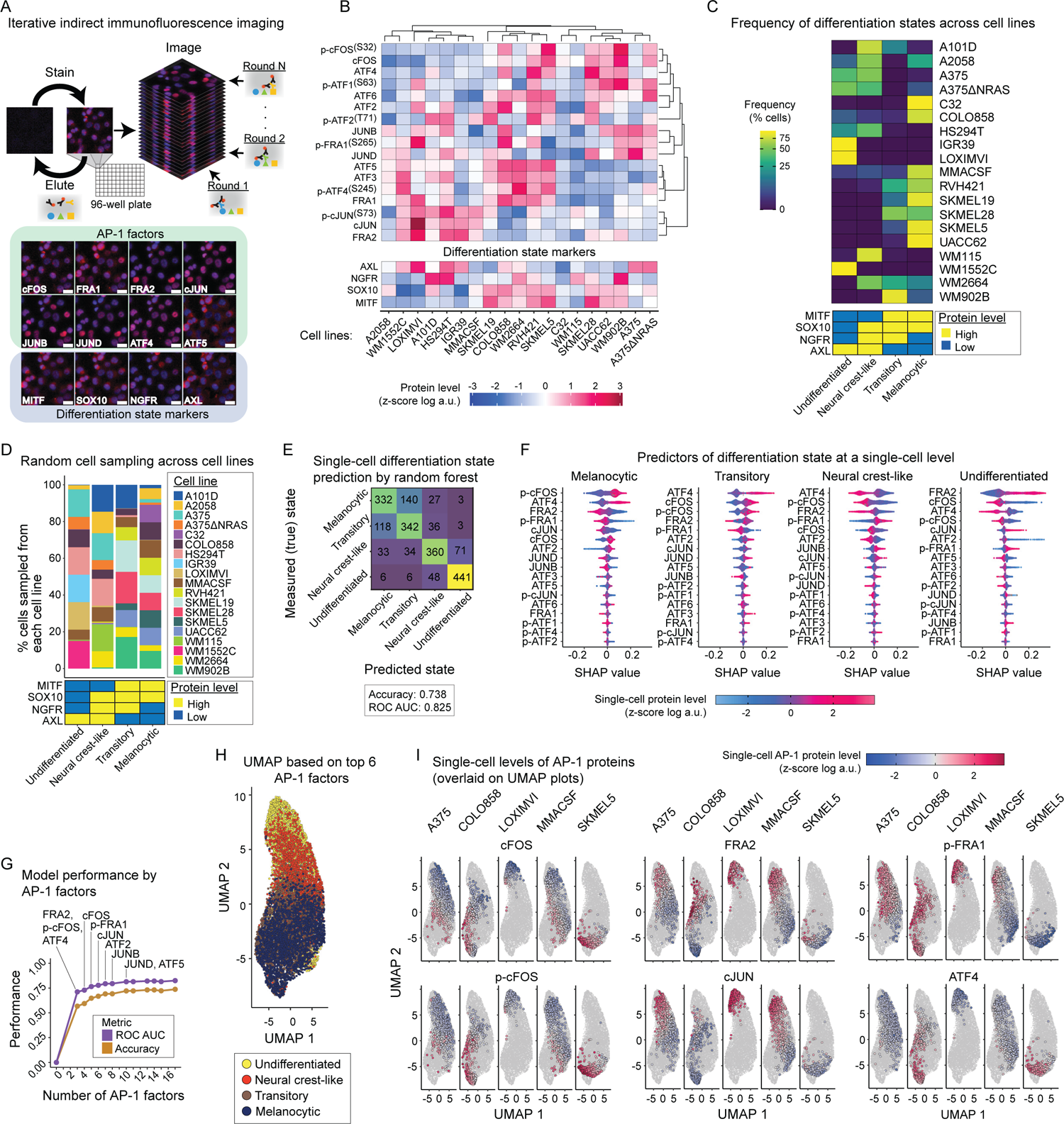
Single-cell AP-1 protein levels predict differentiation state heterogeneity in melanoma cells. **(A)** Schematic representation of the iterative indirect immunofluorescence imaging (4i) procedure used in this study to generate multiplexed single-cell data on 17 AP-1 proteins and 4 differentiation state markers. Representative images of selected AP-1 transcription factors and differentiation state markers are shown for LOXIMVI cells. Scale bars represent 20 µm. Hoechst staining of nuclei is shown in blue while staining of the indicated protein is shown in red. **(B)** Population-averaged measurements of 17 AP-1 proteins and 4 differentiation state markers acquired across 19 BRAF-mutant melanoma cell lines. Protein data shown for each condition represent the log-transformed mean values for two replicates, followed by z-scoring across all cell lines. Data are organized based only on hierarchical clustering of AP-1 protein measurements with Pearson correlation distance metric and the average algorithm for computing distances between clusters. **(C)** Natural frequency of cells in each differentiation state (defined based on MITF, SOX10, NGFR and AXL levels) across 19 BRAF-mutant melanoma cell lines. **(D)** The percentage of cells sampled from each of the 19 cell lines, and their corresponding differentiation states used in the random forest model. **(E)** Confusion matrix showing the independent validation performance of the random forest classifier in predicting the differentiation state of cells based on single-cell AP-1 measurements. The model was trained using a group of 8,000 cells and validated using an independent group of 2,000 cells. The prediction accuracy and area under the receiver operating characteristic curve (ROC AUC) are shown as an overall measure of the classifier performance. **(F)** Distributions of Shapley Additive exPlanations (SHAP) scores for each AP-1 factor across individual cells from the independent validation set. The color indicates the z-score scaled, log-transformed level of each AP-1 protein at a single-cell level. For each differentiation state, AP-1 factors are ordered based on the mean absolute values of their SHAP scores. **(G)** Classification performance of the random forest model based on varying numbers of top AP-1 factors (based on their SHAP values) used as predictors. **(H)** UMAP analysis of the sampled melanoma cells (as shown in **D**) based on their multiplexed levels of top 6 predictive AP-1 measurements (FRA2, p-cFOS, ATF4, cFOS, p-FRA1 and cJUN). Cells are colored based on their differentiation states. **(I)** Single-cell levels of the top six AP-1 proteins overlaid on UMAP plots for representative cell lines.

The population-averaged and single-cell protein data revealed a high degree of variation in differentiation state markers and AP-1 proteins across genetically distinct cell lines (Figures 1B, 1C, S1A). To test whether there is a relationship between AP-1 variations and the differentiation state of individual cells regardless of their genetic differences, we randomly sampled a total of 10,000 cells, including 2,500 from each of the four differentiation states, in a way that they represented all 19 cell lines and 4 distinctive differentiation states as equally as possible (Figure 1D). We used the multiplexed AP-1 data of 80% of the cells to train a random forest classification model to predict the differentiation state of each individual cell. We then used the remaining 20% of the cell population to independently validate model predictions. Model-predicted single-cell differentiation states matched true (measured) differentiation states with an accuracy of 0.74, representing a remarkable performance relative to a random 4-class classifier with an expected accuracy of 0.25 (Figure 1E). A close look at model predictions showed that they matched true states for ∼88% of undifferentiated cells, ∼72% of neural crest-like cells and ∼66% of melanocytic cells. In cases where the true and predicted state of a cell did not match, the model often predicted a closely related neighboring state along the differentiation state trajectory. When we combined cells from these related states, e.g., cells in melanocytic and transitory states, the model was able to distinguish them from the other two states in ∼93% of the cases (Figure 1E).

To identify those AP-1 measurements that most strongly predicted single-cell differentiation state, we computed the SHapley Additive exPlanations (SHAP) values for the random forest classifications (Lundberg et al., 2020). SHAP assigns each AP-1 factor an importance value, quantifying its contribution, either positively or negatively, to the predicted differentiation state of any given cell (Figure 1F). Among the most important AP-1 factors (ranked based on mean absolute SHAP values) were p-cFOS, FRA2, ATF4, cFOS, p-FRA1 and cJUN. Single-cell measurements of these six factors made it possible to predict the differentiation state of a cell with an accuracy of 0.67 (Figure 1G).

We then asked whether models trained based on the top six AP-1 factors would be able to predict the differentiation state of new cells from independent cell lines not included in model training. To answer this question, we iteratively removed one cell line, built a model using randomly sampled cells from the remaining 18 cell lines, and then used the trained model to predict the differentiation state of randomly selected cells from the left-out cell line. We observed that prediction accuracy for left-out cell lines was variable with an average value of 0.49 ± 0.14 across all 19 iterations (Figures S1B, S1C). We also noticed that while prediction accuracy for some left-out cell lines was greater than that of the full model (e.g., ∼0.87 for LOXIMVI cells), predictions for two left-out cell lines (including IGR39 and SKMEL19) underperformed the random model. To test whether the lower performance of model predictions for a few cell lines could be attributed to any common patterns of misclassification, we examined the single-cell predictions for each left-out cell line separately (Figure S1D). We found that in most cases, misclassification occurred when the model failed to distinguish between closely related neighboring states (e.g., neural crest-like versus undifferentiated cells in IGR39, or transitory versus neural crest-like cells in SKMEL19). When we combined cells from such closely related states, the models were able to distinguish them from the other two states in >80% of the cases (Figure S1C).

Together, these analyses revealed that the heterogeneity in melanoma differentiation state was associated with distinguishable patterns of variation in the expression of a few key AP-1 proteins. In agreement with the SHAP analysis and model validation results, dimensionality reduction by Uniform Manifold Approximation and Projection (UMAP) (Becht et al., 2018) using only the top six AP-1 factors resulted in a cell trajectory ordered from melanocytic to undifferentiated states (Figures 1H, 1I). The UMAP projection also shows that melanocytic and transitory cells expressed substantially higher levels of p-cFOS, cFOS and ATF4, while undifferentiated cells exhibited lower levels of all these factors and instead exhibited increased levels of FRA2, cJUN and p-FRA1.

### AP-1 transcript levels predict variations in differentiation state programs across melanoma lines

To test whether the relationships between the patterns of AP-1 expression and melanoma differentiation state were recapitulated at the transcriptional level, we analyzed a previously published dataset, including RNA sequencing of 53 melanoma cell lines (Tsoi et al., 2018). Each cell line was assigned a series of seven signature scores, defined as the average of z-scores for the expression levels of differentiation state signature genes (Tsoi et al., 2018). The differentiation signature scores were then related to the transcript levels of 15 AP-1 genes for each cell line by partial least square regression (PLSR) (Figure 2A). The overall performance of the PLSR model was evaluated by computing the fraction of variance in signature scores explained (R^2^) or predicted (Q^2^) by changes in AP-1 gene expression (Figure 2B). The model revealed a high performance and prediction accuracy with R^2^ of 0.72 and Q^2^ of 0.55 (using leave-one-out cross-validation) for four PLSR components. To evaluate the accuracy of predictions for each differentiation state, we assessed the correlation between the signature scores derived from the differentiation signature genes and scores predicted by the PLSR model. The model showed consistent accuracy with an average Pearson’s correlation coefficient of 0.74 ± 0.08 (*P* = 3.2×10^-17^ to 1.3×10^-6^) between the actual and predicted signature scores (Figure 2A). To independently validate the model predictions, we used RNA sequencing data from a different panel of 32 BRAF-mutant melanoma cell lines in the Cancer Cell Line Encyclopedia (CCLE) (Ghandi et al., 2019). The PLSR model trained against the original set of 53 cell lines was able to predict the differentiation signature scores in the new set of 32 melanoma cell lines, leading to an average Pearson’s correlation coefficient of 0.65 ± 0.13 (*P* = 2.3×10^-8^ to 6.8×10^-3^) between the actual and predicted scores (Figure 2C).

**Figure 2.**
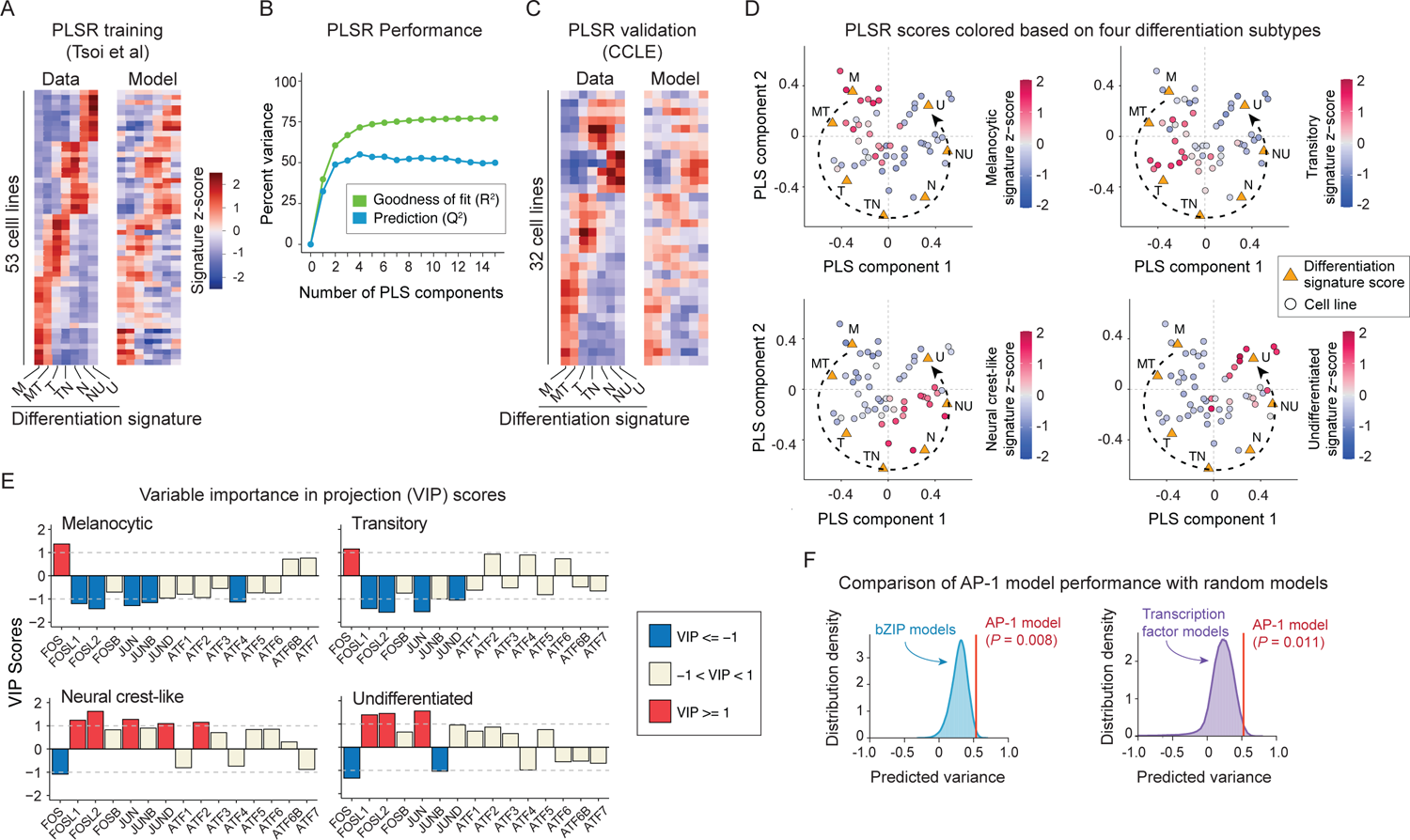
AP-1 transcript levels predict variations in differentiation state across melanoma lines. **(A)** Comparison between differentiation signature scores computed based on RNA sequencing data for 53 cell lines reported by Tsoi *et al* (left) and PLSR-predicted scores (following leave-one-out cross-validation) for each cell line based on their transcript levels of 15 AP-1 genes (right). M: melanocytic; MT: melanocytic-transitory; T: transitory; TN: transitory-neural crest-like; N: neural crest-like; NU: neural crest-like-undifferentiated; U: undifferentiated. **(B)** Performance of the PLSR model evaluated by computing the fraction of variance in differentiation signature scores explained (R^2^) or predicted based on leave-one-out cross validation (Q^2^) with increasing number of PLS components. **(C)** Comparison between differentiation signature scores computed based on RNA sequencing data of 32 CCLE cell lines (left) and predicted scores based on the PLSR model built for the original set of 53 cell lines (right). **(D)** PLSR scores (of the first two PLS components) for each cell line colored according to their differentiation signature scores for melanocytic, transitory, neural crest-like and undifferentiated states. **(E)** PLSR-derived variable importance in projection (VIP) scores, highlighting combinations of AP-1 transcripts that are predictive of differentiation signature scores for melanocytic, transitory, neural crest-like and undifferentiated states. The sign of the VIP score shows whether the indicated variable (AP-1 transcript level) positively or negatively contributes to a given differentiation signature. Significant VIP scores (of greater than 1 or smaller than −1) are highlighted. **(F)** Comparison of performance (with respect to differentiation state prediction) between the PLSR model based on transcript levels of the top 8 AP-1 transcription factors with models based on transcript levels of combinations of 8 randomly chosen bZIP family transcription factors (*n* = 1×10^5^ iterations; left panel) or built based on 8 randomly chosen transcription factors (*n* = 5×10^5^ iterations; right panel). Empirical *P* values were reported for the comparison of predicted variances based on ten-fold cross-validation.

The high performance of the PLSR model shows that variations in the transcriptional levels of at least some AP-1 genes may explain the variability in differentiation states across melanoma cell lines. In agreement with this expectation, different cell lines could be separated by their PLSR scores based on their position along the different state trajectory (Figure 2D). Because the PLSR model achieved its maximum prediction accuracy by four components, we computed the Variable Importance in the Projection (VIP) scores across all these components to determine the overall contribution of each AP-1 gene to each differentiation state (Figure 2E). Among the most important predictors of differentiation state (determined by |VIP| > 1) were the expression of FOS (encoding cFOS), FOSL1 (encoding FRA1), FOSL2 (encoding FRA2), JUN (encoding cJUN), JUNB, JUND, ATF2 and ATF4 (Figure 2E). Importantly, a model created using only these AP-1 genes was able to significantly outperform most PLSR models that were built based on combinations of eight randomly chosen transcription factors from the basic leucine zipper (bZIP) family (the family to which AP-1 factors belong) (*P* = 0.008) or based on any eight randomly chosen transcription factors (*P* = 0.01) (Figure 2F).

Together, these analyses revealed that the predictivity of patterns of AP-1 variation for melanoma differentiation state could also be captured at the level of transcription of these factors. Except for ATF4, the statistical association of AP-1 factors with differentiation state was generally consistent across bulk transcript and single-cell protein measurements (Figure S2A). Melanocytic and transitory cells expressed substantially higher levels of FOS transcript and cFOS protein levels, whereas undifferentiated cells were associated with increased levels of FOSL1, FOSL2 and JUN transcripts and their corresponding proteins FRA1, FRA2 and cJUN, respectively.

### Single-cell network inference reveals the role of AP-1 activity in regulation of differentiation programs

Next, we asked whether the statistical associations between the identified key AP-1 proteins and single-cell differentiation states resulted from the active regulation of differentiation programs by the AP-1 factors. To address this question, we applied single-cell regulatory network inference and clustering (SCENIC) (Aibar et al., 2017; Van de Sande et al., 2020) to analyze a previously published single-cell RNA sequencing dataset of 10 melanoma cell lines (Wouters et al., 2020). SCENIC uses single-cell gene expression data to infer transcription factors alongside their candidate target genes (collectively called a regulon), enabling the identification of regulatory interactions and transcription factor activities with high confidence. In line with our results from the gene and protein expression analyses, SCENIC analysis found the FOSL2 and JUN motif regulons to be substantially enriched in populations of undifferentiated cells in comparison with melanocytic, transitory, and neural crest-like cells (Figures 3A, 3B). The activity of the FOSL1 regulon was low in melanocytic cells but gradually increased among transitory, neural crest-like and undifferentiated cells (Figure 3C). The FOS regulon, on the other hand, was substantially enriched in melanocytic and transitory cells, but its activity was low in undifferentiated and neural crest-like cells (Figure 3D).

**Figure 3.**
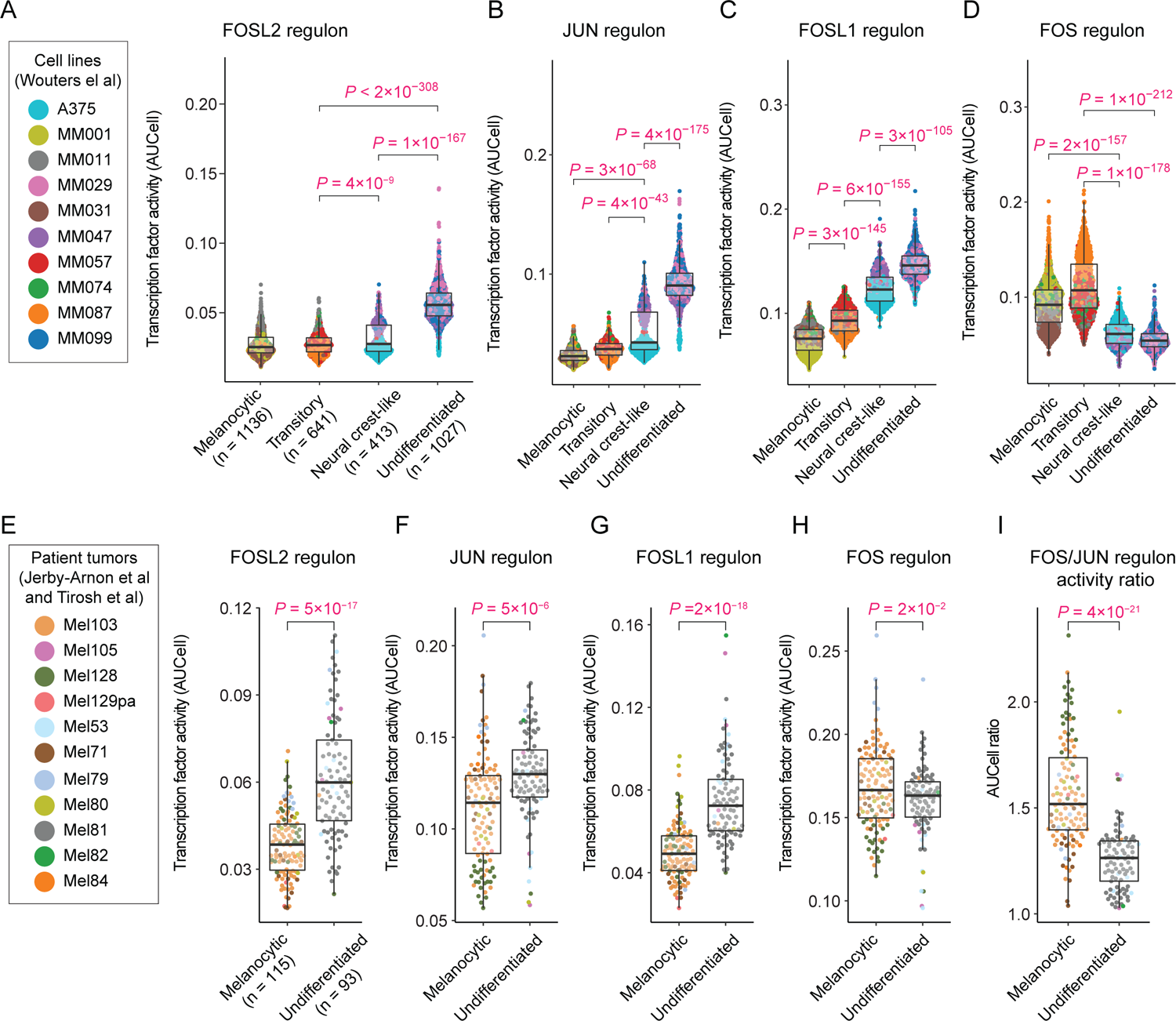
Single-cell network inference reveals the role of AP-1 activity in regulation of differentiation state programs. **(A-D)** Single-cell distributions of the activity of SCENIC regulons for FOSL2 **(A)**, JUN **(B)**, FOSL1 **(C)** and FOS **(D)** motifs, measured using AUCell in individual cells (from 10 melanoma cell lines profiled by Wouters *et al*) across distinct differentiation states. The differentiation state of individual cells was determined based on their gated levels of enrichment (quantified by AUCell) for the differentiation gene signatures as defined by Tsoi *et al*. **(E-I)** Single-cell distributions of the AUCell activity of SCENIC regulons for FOSL2 **(E)**, JUN **(F)**, FOSL1 **(G)**, and FOS **(H)** motifs, as well as the ratio of FOS and JUN regulon activities **(I)**, quantified in individual cells from 11 treatment-naïve melanoma tumors as profiled by Tirosh *et al* and Jerby-Arnon *et al*. Statistical comparisons were performed using two-sided unpaired *t* test. Boxplot hinges correspond to the lower and upper quartiles, with a band at median. Whiskers indicate 1.5 times interquartile ranges.

To test whether the relationship between AP-1 regulon activities and melanoma differentiation states existed in single cells derived from tumor biopsies, we performed differentiation state enrichment and SCENIC analysis on single-cell RNA sequencing data previously collected via dissociation and profiling of patient-derived melanoma samples (Jerby-Arnon et al., 2018; Tirosh et al., 2016).

Accounting for missing values in a subset of differentiation signature genes, enrichment analysis of the 11 treatment-naïve melanoma samples distinguished single cells from melanocytic and undifferentiated states with high confidence. SCENIC analysis of these cells showed that FOSL2, JUN and FOSL1 regulons were significantly enriched in undifferentiated cells in comparison with melanocytic cells (Figures 3E-3G). In contrast to a substantially higher FOS regulon activity observed in cultured melanocytic cells, the activity of FOS regulon was only slightly higher in melanocytic tumor cells in comparison with undifferentiated cells (Figure 3H). Interestingly, however, the FOS/JUN activity ratio at a single-cell level was able to distinguish melanocytic cells from undifferentiated cells more efficiently than either of these AP-1 factors alone (Figure 3I), suggesting that it is the balance between AP-1 factor activities that determines a cell’s differentiation state.

Next, we asked how the key AP-1 transcription factor activities and their activity ratios varied between the widely recognized two-class “proliferative” and “invasive” phenotypes in melanoma cells (Hoek et al., 2006). Following single-cell enrichment analysis of the transcriptional signatures defined for these phenotypes (Hoek et al., 2006), we compared their associations with the activity of key AP-1 regulons inferred by SCENIC. As expected, the invasive phenotype exhibited higher activities of FOSL1, FOSL2 and JUN, whereas proliferative cells showed a higher FOS/JUN activity ratio (Figure S2B-S2K).

Together, these analyses revealed that melanoma cells of diverse differentiation states are associated with distinct regulatory network activities by AP-1 transcription factors. In particular, the role of FOS, FOSL1, FOSL2 and JUN regulon activities was consistent with their corresponding patterns of gene and protein expression across melanoma differentiation states at both population and single-cell levels.

### MAPK inhibitor-induced changes in the AP-1 state predict changes in differentiation state

Although melanoma populations consist of stable mixtures of cells in diverse differentiation states at baseline, they can switch state in response to environmental perturbations. Specifically, treatment with MAPK inhibitors has been reported to induce changes in cell state that are associated with either activation of an MITF^High^ program triggering melanocytic differentiation (Rambow et al., 2018; Smith et al., 2016), or downregulation of MITF activity and induction of an NGFR^High^, neural crest-like state (Fallahi Sichani et al., 2017; Rambow et al., 2018). Such adaptive phenotype switching occurs with as little as 3 days of exposure to MAPK inhibitors and concomitantly with changes in MITF and NGFR protein expression (Khaliq et al., 2021). To determine common patterns of AP-1 changes that might be associated with drug-induced changes in differentiation state in either direction (i.e., differentiation or dedifferentiation), we exposed 18 BRAF-mutant melanoma cell lines to the BRAF inhibitor vemurafenib (at 0.316 µM) either alone or in combination with the MEK inhibitor trametinib (at 0.0316 µM). We fixed the cells following 24 or 72 h of treatment and then used the 4i procedure to measure the abundance or phosphorylation state of AP-1 transcription factors as wells as MITF and NGFR protein levels (Figures 4A, S3, S4, S5A). We also measured p-ERK^T202/Y204^ levels to quantify changes in MAPK signaling as described in the following section (Figure S5B).

**Figure 4.**
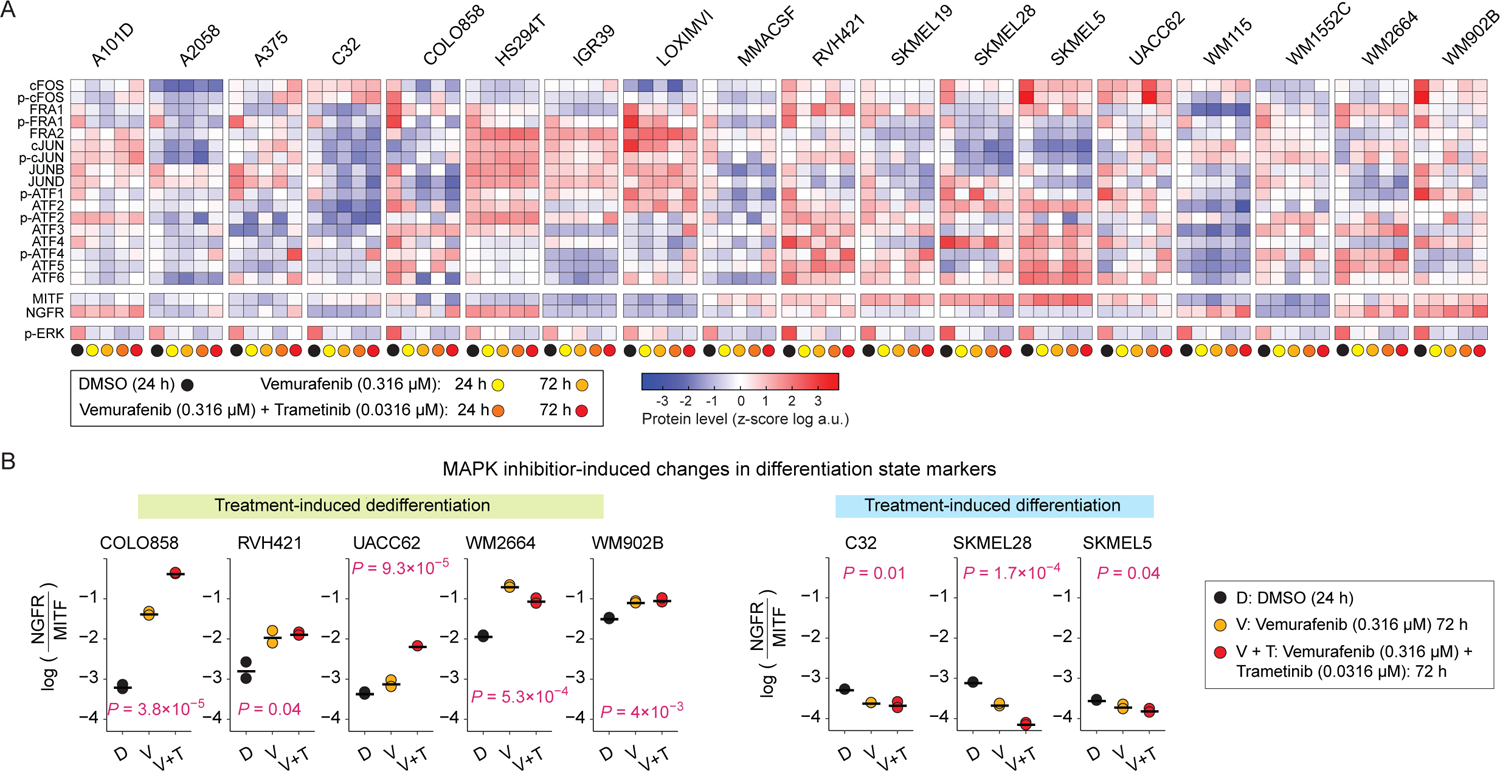
MAPK inhibitor-induced changes in AP-1 protein levels, p-ERK and differentiation state markers. **(A)** Population-averaged measurements of 17 AP-1 proteins, differentiation state markers MITF and NGFR, and p-ERK^T202/Y204^ levels acquired across 18 BRAF-mutant melanoma cell lines. Protein data shown for each condition represent the log-transformed mean values for two replicates, followed by z-scoring across all cell lines and treatment conditions, including DMSO, vemurafenib alone (at 0.316 µM) or the combination of vemurafenib (at 0.316 µM) and trametinib (at 0.0316 µM) for 24 or 72 h. **(B)** MAPK inhibitor-induced changes in differentiation state, as evaluated by log-transformed ratio of NGFR to MITF protein levels across cell lines at 72 h. Central marks on the data points indicate the mean between two replicates*. P-*values represent one-way ANOVA test for differences across treatment conditions in each cell line.

To assess drug-induced changes in differentiation state for each cell line, we computed the relative enrichment of dedifferentiated cells by normalizing the NGFR protein levels to MITF protein levels at baseline (DMSO), then tracking its changes following MAPK inhibitor treatments. Interestingly, treatment with BRAF/MEK inhibitors induced dedifferentiation in some cell lines (Figure 4B; left panels) but enhanced differentiation in a few others (Figure 4B; right panels). To identify possible associations between AP-1 factors and drug-induced changes in differentiation state, we built a PLSR model to associate DMSO-normalized changes in the expression levels of each of the AP-1 factors to DMSO-normalized changes in the enrichment of dedifferentiated (or abatement of differentiated) cells for each of the MAPK inhibitor treatment conditions. The PLSR model achieved its maximum prediction accuracy by three components (Figure 5A). We thus computed the VIP scores using the first three PLS components to determine the overall contribution of each AP-1 modification to drug-induced changes in melanoma differentiation state (Figure 5B). VIP analysis revealed changes in multiple AP-1 factors from the JUN and ATF subfamily that were correlated with changes in differentiation state. Among these factors, cJUN and p-cJUN changes (at 24 h and 72 h) were consistently the strongest predictors of both drug-induced dedifferentiation and differentiation among the tested cell lines (Figure 5C). In agreement with population-level data, single-cell analysis revealed increases in cJUN and p-cJUN levels in dedifferentiating melanoma cells (Figure 5D, top row), and reduction of cJUN and p-cJUN levels in differentiating melanoma cells following MAPK inhibitor treatments (Figure 5D, bottom row).

**Figure 5.**
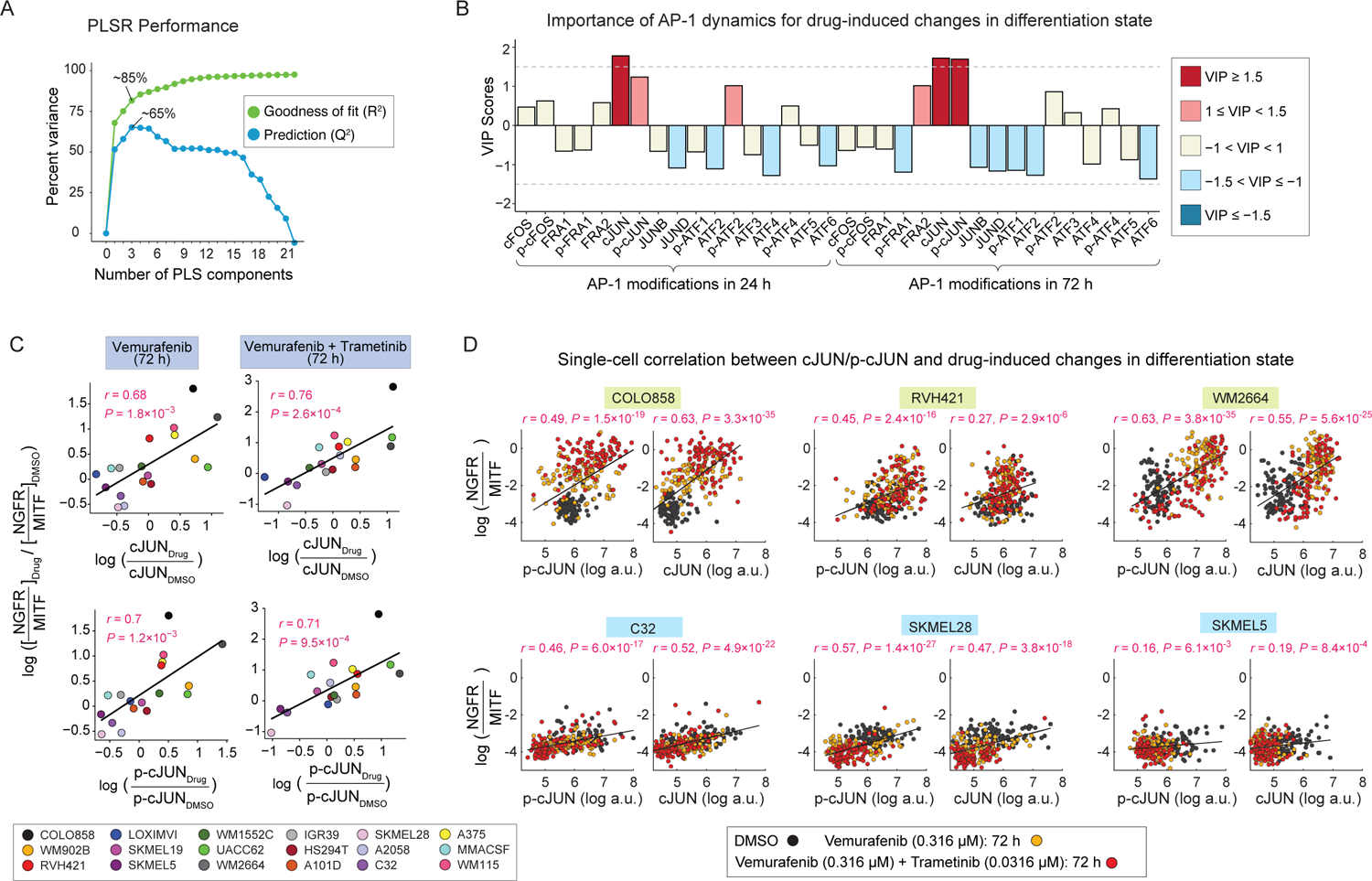
MAPK inhibitor-induced changes in cJUN and p-cJUN levels correlate with drug-induced changes in differentiation states. **(A)** Performance of the PLSR model in predicting drug-induced changes in differentiation states based on drug-induced AP-1 modifications. Model performance was evaluated by computing the fraction of variance in DMSO-normalized differentiation state changes at 72 h explained (R^2^) or predicted based on leave-one-out cross validation (Q^2^) with increasing number of PLS components. **(B)** PLSR-derived variable importance in projection (VIP) scores, highlighting combinations of DMSO-normalized AP-1 modifications at 24 and 72 h, and their importance for predicting the DMSO-normalized differentiation state changes at 72 h. The sign of the VIP score shows whether the indicated variable (DMSO-normalized AP-1 protein levels at 24 h or 72 h) positively or negatively contributes to the response (DMSO-normalized change in differentiation state). **(C)** Pearson’s correlation between DMSO-normalized changes in differentiation state and DMSO-normalized cJUN or p-cJUN levels at indicated timepoints and drug treatment conditions. Each data-point represents population-averaged measurements across two replicates for each cell line. **(D)** Analysis of covariance between the levels of p-cJUN or c-JUN and drug-induced changes in differentiation state (as evaluated by NGFR-MITF ratio) at the single-cell level across indicated treatment conditions. For each cell line, Pearson’s correlation coefficient was calculated based on 300 randomly sampled cells (100 cells from each treatment condition).

### MAPK inhibitor-induced changes in the AP-1 state reveal efficiency of ERK pathway inhibition across melanoma cell lines

In addition to drug-induced changes in differentiation state, incomplete inhibition of the ERK pathway (or its reactivation following a transient period of ERK inhibition) is known as a common mechanism of resistance to BRAF inhibitors (Lito et al., 2012). Importantly, when ERK activity rebounds as early as a few hours after drug treatment, a small residual ERK activity (at the cell population level) may be sufficient to help cells escape the effect of BRAF inhibition (Gerosa et al., 2020; Khaliq et al., 2021). The combination of BRAF inhibitors with MEK inhibitors has been proposed as a strategy to overcome the ERK pathway rebound following BRAF inhibition alone (Lito et al., 2012). In agreement with this idea, all 18 melanoma cell lines treated with the combination of vemurafenib and trametinib showed significantly lower levels of residual p-ERK at 24 h in comparison with their responses to vemurafenib treatment for the same duration (Figure 6A, B). Drug combination also significantly reduced p-ERK rebound in comparison with vemurafenib treatment at 72 h. However, the extent of this effect was variable among different cell lines (Figure 6A, B). We thus asked which, if any, of the AP-1 factors might capture the observed differences in the efficiency of ERK pathway inhibition among different cell lines.

**Figure 6.**
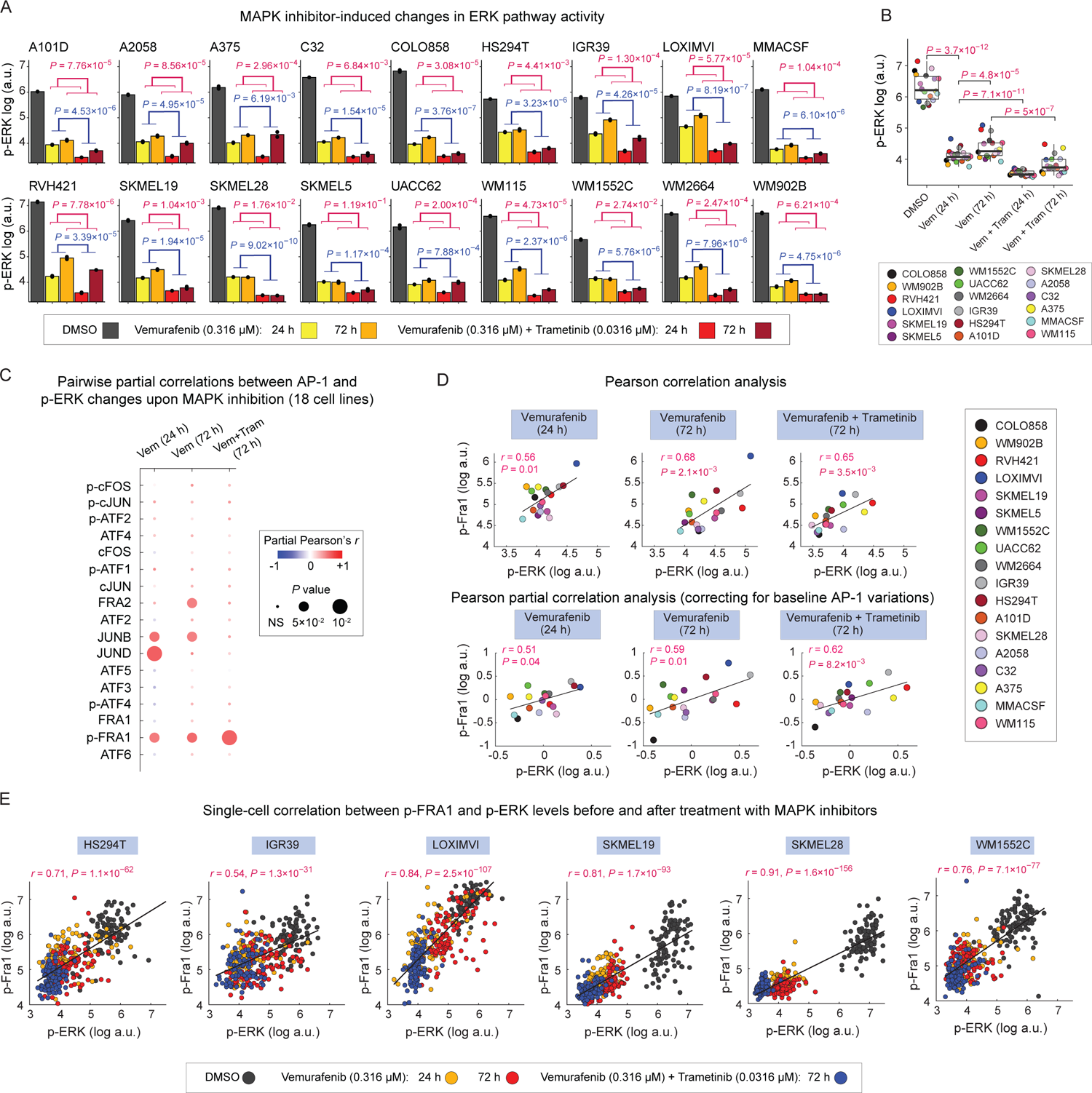
MAPK inhibitor-induced changes in p-FRA1 levels correlate with efficiency of ERK pathway inhibition across cell lines. **(A)** Population-averaged measurements of p-ERK^T202/Y204^ levels in 18 cell lines following indicated MAPK inhibitor treatments for 24 h or 72 h. Bar height indicates mean values between two replicates shown as black dots. *P*-values show the statistical significance of the impact of MAPK inhibitor treatment (blue) or time (red) on p-ERK levels based on two-way ANOVA. **(B)** Statistical comparison (using two-sided paired *t* test) of p-ERK levels across 18 cell lines following indicated MAPK inhibitor treatments for 24 h or 72 h. Each data-point represents population mean of p-ERK levels between two replicates for each cell line. **(C)** Pairwise partial correlations (evaluated across 18 cell lines) between each of the 17 AP-1 measurements and p-ERK levels following 24 or 72 h of treatment with MAPK inhibitors, while correcting for the corresponding baseline (drug-naïve) AP-1 levels in the same cell lines. **(D)** Pearson’s correlation (top row) and partial correlation (bottom row) between p-ERK and p-FRA1 levels following MAPK inhibitor treatments at indicated timepoints. Each data-point represents population mean of p-ERK levels between two replicates for each cell line. **(E)** Analysis of covariance between p-FRA1 and p-ERK levels across indicated MAPK inhibitor treatment conditions at the single-cell level. For each cell line, Pearson’s correlation coefficient was calculated based on 400 randomly sampled cells (100 cells from each treatment condition).

To answer this question, we used partial correlation analysis to assess pairwise relationships between p-ERK and AP-1 levels across 18 cell lines treated with either vemurafenib (for either 24 or 72 h) or the combination of vemurafenib and trametinib (for 72 h), while correcting for baseline (drug-naïve) variations in the AP-1 protein levels (Fig. 6C). This analysis identified p-FRA1 as the most consistent predictor of the efficiency of ERK pathway inhibition among all cell lines (Fig. 6C, 6D). In agreement with population-level correlation analysis, single-cell analysis also revealed a significant covariance between p-ERK and p-FRA1 levels (Figure 6E). Such strong connection between p-FRA1 and p-ERK in drug-treated cells is consistent with FRA1 serving as a tightly coupled sensor of ERK activity (Gillies et al., 2017). Interestingly, however, FRA1 or p-FRA1 levels did not correlate with hyperactivated p-ERK levels when we performed pairwise correlation analysis on drug-naïve BRAF-mutant cells. Instead, we found drug-naïve p-ERK levels to be positively correlated with cFOS, p-cFOS and ATF4, and negatively correlated with FRA2 and cJUN (Figure S6). All these AP-1 factors were predictors of melanoma differentiation state. These observations are consistent with previous reports linking up-regulation of MITF to elevated ERK activity in BRAF-mutant melanoma cells (Wellbrock et al., 2008). In addition, they suggest that changes in ERK signaling following pharmacological inhibition of the pathway may lead to rewiring of AP-1 signaling.

### Perturbation of AP-1 state by siRNA confirms its role in driving differentiation state heterogeneity

We hypothesized that if the AP-1 state drives melanoma differentiation programs, then inducing perturbations in the AP-1 state will shift differentiation states in predictable ways. To test this hypothesis, we perturbed AP-1 factors in COLO858 melanoma cells by using pools of previously validated siRNAs to knock down the expression of five AP-1 genes, including FOS, FOSL1, FOSL2, JUN, and JUND, either individually or in pairwise combinations. Unfortunately, we were unable to achieve significant depletion of FRA1 in COLO858 cells. Hence, we focused on the analysis of the data following individual, or pairwise combinations of siRNAs targeting FOS, FOSL2, JUN, and JUND. COLO858 represents a heterogenous population composed of both melanocytic (SOX10^High^/ MITF^High^) and undifferentiated (SOX10^Low^/ MITF^Low^) cells, thereby allowing us to track changes in the expression of differentiation state markers after AP-1 perturbations. Following 96 h of AP-1 gene knockdown in COLO858 cells, we measured (in three replicates) protein levels of differentiation markers MITF, SOX10 and AP-1 factors cFOS, FRA1, FRA2, cJUN, and JUND using 4i (Figures 7A, 7B, S7) Interestingly, siRNA-mediated knockdowns not only reduced the levels of AP-1 proteins targeted by their corresponding siRNAs but also, in some cases, led to changes in the expression of other AP-1 proteins (Figures 7A, S7). For example, FOSL2 knockdown substantially reduced FRA2 levels, but also induced the expression of FRA1, when compared to cells treated with non-targeting (control) siRNA. JUND knockdown reduced JUND levels, but also led to an increase in cFOS and cJUN levels. These observations agree with previous reports (Lopez-Bergami et al., 2010), suggesting the state of AP-1 network is controlled by interactions among different AP-1 factors. Combinations of siRNAs against pairs of AP-1 genes may help reveal AP-1 modifications that are phenotypically consequential. To identify such interactions, we quantified the levels of SOX10 and MITF proteins across all knockdown conditions in COLO858 cells and examined them along a two-dimensional plot (Figures 7A, 7B). We found that knocking down FOSL2 and JUN in combination significantly increased the expression of SOX10 (Figures 7A, 7B). This behavior is consistent with our earlier finding regarding the role of FRA2 and cJUN in regulation of the undifferentiated (SOX10^Low^) state. On the other hand, knocking down FOS and JUND in combination significantly reduced the expression of both MITF and SOX10 (Figures 7A, 7B), which is consistent with our finding regarding their role in driving the melanocytic lineage.

**Figure 7.**
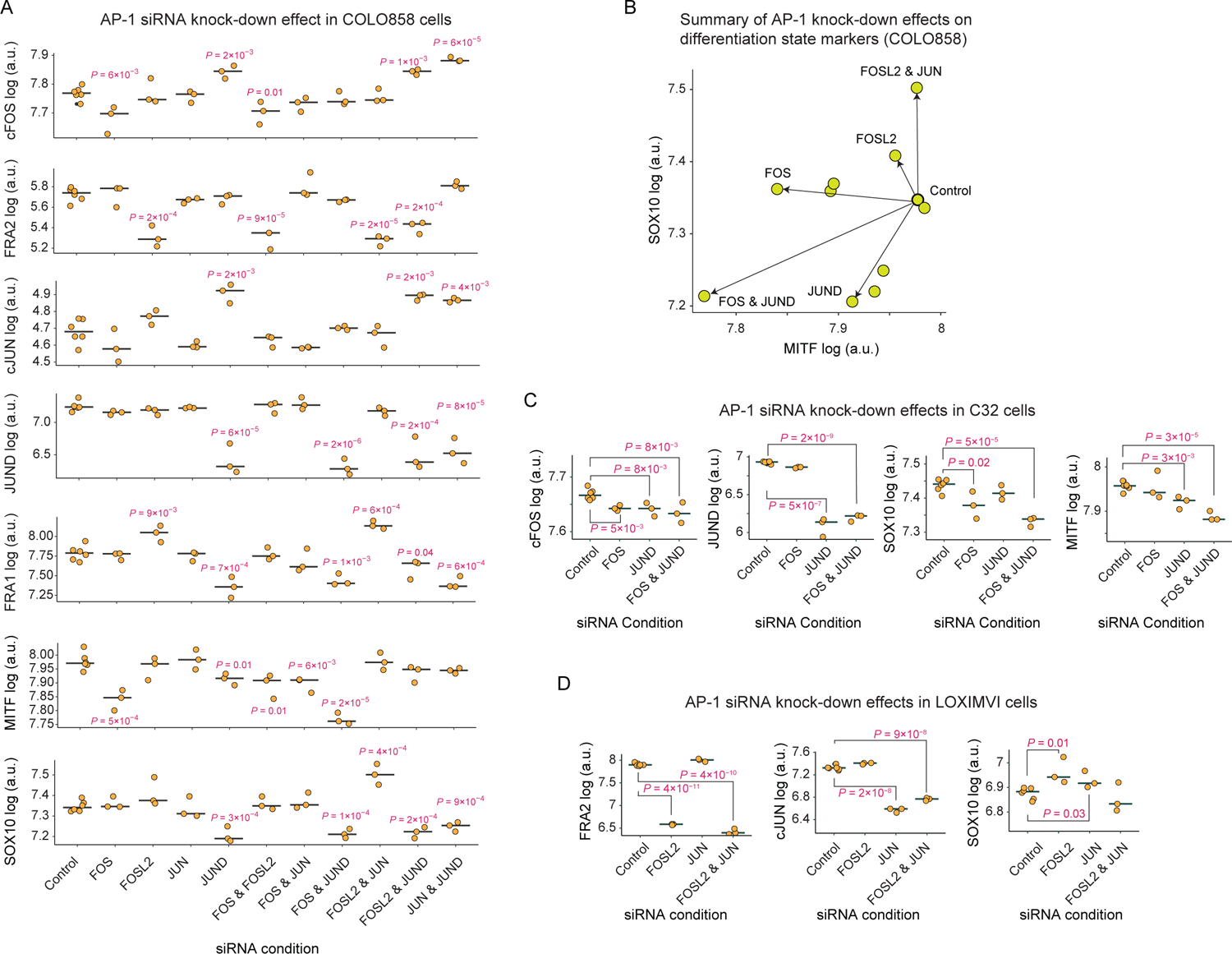
Perturbation of AP-1 state by siRNA confirms its role in driving differentiation state heterogeneity. **(A)** The effect of siRNA-mediated depletion (for 96 h) of AP-1 proteins cFOS, FRA2, cJUN, and JUND, either individually or in pairwise combinations, on protein levels of cFOS, FRA1, FRA2, cJUN, JUND and differentiation state markers MITF and SOX10 in COLO858 cells. Protein data shown for each condition represent the log-transformed mean values for three and six replicates across AP-1 knockdown conditions and the control, respectively. The central mark on the plots indicates the median across replicates. Statistical comparisons were performed using two-sided unpaired *t* test. **(B)** Two-dimensional projection of MITF and SOX10 levels (in log scale) following 96 h of siRNA knockdown in COLO858 cells. **(C, D)** Statistical comparison (using two-sided unpaired *t* test) of selected AP-1 proteins and SOX10 levels across indicated siRNA knockdown conditions in C32 **(C)** and LOXIMVI **(D)** cells. Protein data shown for each condition represent the log-transformed mean values for three and six replicates across AP-1 knockdown conditions and the control, respectively. The central mark on the plots indicates the median across replicates.

To further validate the impact of depletion of AP-1 proteins on the expression of melanoma differentiation state markers, we performed selected siRNA knockdown experiments in two additional cell lines, including C32 and LOXIMVI which constituted relatively homogeneous populations of melanocytic and undifferentiated cells, respectively (Figure 1C). First, we exposed C32 cells to siRNAs targeting FOS and JUND, individually or in combination for 96 h (Figure 7C). In agreement with results in COLO858 cells, the combination of FOS and JUND knockdown significantly reduced SOX10 and MITF protein levels in C32 cells (Figure 7C). Next, we exposed undifferentiated LOXIMVI cells to siRNAs targeting FOSL2 and JUN. We found that both siRNAs individually increased the expression levels of SOX10 (Figure 7D). However, in contrast to our observation in COLO858 cells, the impact of combined FOSL2 and JUN siRNAs (for 96 h) in highly undifferentiated LOXIMVI cells was not significant. Overall, despite some minor differences among the impact of individual or combined siRNA treatments, the knockdown experiments across three cell lines (taken with all the data presented throughout this study) confirmed our findings regarding a key role for a balance between FOS, JUND, JUN and FOSL2 in driving the differentiation program in melanoma cells.

## Discussion

The hyperactivation of MAPK signaling in BRAF-mutant melanomas is linked to their overall sensitivity to MAPK inhibition. The differentiation state heterogeneity, however, leads to variability in MAPK inhibitor responses both across genetically diverse tumors and among genetically homogeneous populations of cells. Understanding the origins of such heterogeneity is key to identifying effective strategies to overcome fractional responses that undermine the potential of MAPK-targeted therapies. It requires a detailed knowledge of the mechanisms and molecular players that link epigenetic plasticity and transcriptional regulation of differentiation state to therapy-induced changes in MAPK signaling. To begin to fill this gap in our knowledge, we used a multidimensional approach at single-cell resolution to systematically investigate the AP-1 transcription factor contributions to heterogeneity in BRAF-mutant melanoma cells. We focus on the AP-1 factors because they serve as downstream targets of MAP kinases, and previous work has connected several AP-1 proteins to MAPK inhibitor resistance, differentiation state heterogeneity, and therapy-induced changes in differentiation state in melanomas.

Our data showed that a tightly regulated balance between a few key AP-1 family members and their activities strongly predict previously characterized heterogeneities in melanoma differentiation states. Specifically, cFOS and p-cFOS were associated with melanocytic and transitory cells, while FRA1, p-FRA1, FRA2, and cJUN and p-cJUN correlated with less differentiated cell states. The systematic nature of the study across many genetically different melanomas suggests that these associations are a general feature of melanomas and likely not unique to a particular cell line or linked to a certain genetic context. Furthermore, we showed that perturbing the molecular balance of AP-1 factors in melanoma cells by siRNAs that deplete specific AP-1 proteins, either alone or in combination, or by treatments with MAPK inhibitors can induce differentiation state switching and heterogeneity in a controllable manner. Together, these findings provide new insights into AP-1 function, its role in cell state plasticity, and its potential dysregulation in melanoma, while opening new avenues for interrogating the AP-1 behavior in the context of adaptive response to MAPK inhibitors. In theory, gaining the ability to target certain AP-1 states could force cells to remain in a more drug-sensitive state, thereby increasing the fractional killing of melanoma cells in response to MAPK inhibition.

Future studies may leverage the findings from this study to further elucidate transcriptional mechanisms that contribute to MAPK-targeted therapy escape in melanomas at a single-cell level. Furthermore, uncovering how the information encoded in the MAPK signaling dynamics is transduced through its downstream AP-1 network will be key for explaining the observed variability in tumor cell responses to MAPK inhibitors. For example, AP-1 family members FOS and FOSL1 are early ERK target genes whose regulation by ERK activity constitutes feedforward motifs that enable them to decode the dynamics of ERK signaling (Davies et al., 2020). Differentiation state-specific variations in the baseline expression and activities of these AP-1 genes, as we observed in this study, could introduce variability in the transduction of MAPK signals, generating heterogeneity in cell fate under MAPK perturbations. Future studies that link dynamic fluctuations in ERK activity and other MAP kinases to AP-1 behavior could offer important insights into mechanisms of heterogeneity in drug response and adaptive resistance to MAPK inhibitors.

Consistent with our findings regarding the role of AP-1 state changes in determining the differentiation state plasticity, recent studies have highlighted a key role for AP-1 factors in chromatin organization and enhancer accessibility. AP-1 proteins have been reported to facilitate new cell fate transitions, such as cellular senescence or differentiation, by establishing the enhancer landscape and granting long-term chromatin access to other transcription factors, thereby allowing the timely execution of cell state-specific transcriptional programs (Madrigal and Alasoo, 2018; Martínez-Zamudio et al., 2020; Phanstiel et al., 2017; Vierbuchen et al., 2017). Understanding which AP-1 factors and cofactors work to keep poised enhancers accessible and which function to shift enhancers from a poised to active state could connect transcription to (de)differentiation and genome reorganization following inhibition of MAPK signaling and its adaptive reactivation. Furthermore, AP-1 proteins like other bZIP proteins must form dimers before they could bind to the AP-1 motif site. For example, while FOS family members bind DNA as obligate heterodimers with members of the JUN family, JUN family members can bind the AP-1 motif site as both homodimers and heterodimers with FOS family members. Future studies, therefore, should also determine the extent to which the combinatorial activity of AP-1 family members is influenced by distinct patterns of dimerization among these transcription factors.

### Limitations of the study

While quantitative, immunofluorescence-based measurements of protein levels and phenotypes and the use of siRNAs to knock down the expression of genes have proven immensely powerful for the study of biology, all techniques have limitations. For example, the quality of quantitative information retrieved from immunofluorescence images depends largely on the quality of segmentation of the cell features of interest. While great effort was made to optimize the image segmentation procedure and to ensure that the segmentation captured the desired features, it is infeasible to visually inspect every cell for appropriate segmentation. Second, in the current manuscript we assume that AP-1 protein levels in the nucleus correspond to or correlate with transcriptional activity. While this assumption is likely true in many circumstances, it is impossible to definitively prove this is the case for each of the many factors we measure in this study. Furthermore, when performing many sequential rounds of 4i staining and images, cells are gradually lost over the course of sample washing and re-probing. While we assume that cell loss occurs equivalently for all differentiation states of the cell, we do not explicitly test this assumption here. Though if this assumption were false, it would be unclear whether the conclusions reached in the paper would change, since our findings are validated using a variety of complementary methods and independent datasets.

## Acknowledgments

We thank Kevin Janes and members of the Fallahi-Sichani laboratory for helpful suggestions and discussion. This work was supported by NIH grant R35-GM133404 (to MFS), the Farrow Fellowship (to DB) and P30-CA044579 (University of Virginia Cancer Center Support Grant). We acknowledge Research Computing at The University of Virginia for providing computational resources and technical support.

## Author Contributions

N.C.-L., D.G.B. and M.F.-S. conceived and designed the study and wrote the manuscript; N.C.-L. performed computational model analysis. D.G.B. developed protocols and performed the experiments. N.C.-L. and D.G.B. performed experimental data analysis. M.F.-S. supervised the work.

## STAR Methods

### RESOURCE AVAILABILITY

#### Lead contact

Further information and requests for resources should be directed to and will be fulfilled by the lead contact, Mohammad Fallahi-Sichani.

#### Materials availability

This study did not generate new unique reagents.

#### Data and code availability

All analyzed data reported in this study are included in the supplemental information. Accession numbers or links for publicly available gene expression and regulon datasets analyzed in this study are listed in the key resources table. Unprocessed data reported in this paper will be shared by the lead contact upon request. All data analysis codes have been deposited at Github and are publicly available as of the date of publication. The link to the Github repository is listed in the key resources table. Any additional information required to reproduce this work is available from the lead contact upon request.

### EXPERIMENTAL MODEL AND SUBJECT DETAILS

#### Cell lines and tissue culture

BRAF-mutant melanoma cell lines used in this study were obtained from the following sources: COLO858, RVH421, A375, A375(NRAS^Q61K^), C32, A2058, WM115, SKMEL28, HS294T, WM1552C, SKMEL5, A101D, and IGR39, LOXIMV1, MMACSF, WM902B and WM2664, UACC62 and SKMEL19 (from the Cancer Cell Line Encyclopedia). All cell lines have been subjected to re-confirmation by short tandem repeat (STR) profiling by ATCC and mycoplasma testing by MycoAlert™ PLUS Mycoplasma Detection Kit. A375, A375(NRAS^Q61K^), A2058, HS294T, A101D, and IGR39 cells were grown in DMEM with 4.5 g/l glucose supplemented with 5% fetal bovine serum (FBS). SKMEL5 and WM2664 cells were grown in EMEM supplemented with 5% FBS. C32, MMACSF, SKMEL28, and WM115 cells were grown in DMEM/F12 supplemented with 1% sodium pyruvate and 5% FBS. COLO858, LOXIMVI, RVH421, SKMEL19, UACC62, WM1552C, and WM902B cells were grown in RPMI 1640 supplemented with 1% sodium pyruvate and 5% FBS. Cells were grown at 37°C with 5% CO_2_ in a humidified chamber. 100 U/mL Penicillin-Streptomycin (10,000 U/mL), and 0.5 mg/mL Plasmocin Prophylactic were present in all cell cultures.

## METHOD DETAILS

### Drug treatments

Cells were seeded in 200 µl/well in Corning 96-well plates. Vemurafenib, Trametinib or vehicle (DMSO) was added at indicated concentrations using the Tecan D300e Digital Dispenser 24 hours after cell seeding. Cells were fixed at the indicated timepoints with 4% paraformaldehyde in phosphate-buffered saline (PBS) for 30 minutes at room temperature. All time course experiments with drug treatment were initiated at the same time and then stopped sequentially at the indicated timepoints (24 h and 72 h). The DMSO experiments were stopped at 24 h to avoid artifacts in signaling measurements that may arise due to cell confluency and the exhaustion of growth media.

### AP-1 gene knockdown by siRNA

COLO858 cells were seeded in 100 µl of antibiotic-free growth media (RPMI supplemented with 5% FBS and 1 mM Sodium Pyruvate) in 96-well plates at a density of 2000 cells/well. After 24 h of incubation, cells were transfected using 0.05 µl of DharmaFECT 2 reagent per well with indicated Dharmacon ON-TARGETplus AP-1 siRNAs (at 25 nM) individually or in pairwise combinations. Knockdowns targeting a single AP-1 gene were supplemented with non-targeting siRNA to normalize the final siRNA concentration (to 50 nM siRNA) across all siRNA conditions. All siRNAs were tested for knockdown efficiency and specificity by measuring protein levels of each factor and members of the factor subfamily (i.e., measure single-cell protein levels of cFOS, FRA1, and FRA2 following FOS knockdown). Only siRNA species that showed knockdown of the protein target without off-target knockdown effects were used. Cells were fixed 96 hours after transfection with 4% paraformaldehyde in PBS for 30 minutes at room temperature. The siRNA sequences used for each condition are included in Table S1.

### Iterative indirect immunofluorescence imaging (4i)

4i images were obtained using a previously described protocol (Gut et al., 2018) with minor modifications. After media aspiration, cells in 96-well plates were fixed with 4% paraformaldehyde in PBS for 30 minutes at room temperature. All washes were performed using a BioTek EL406 Washer Dispenser and consisted of 4 wash cycles of 200 µl with the indicated buffer while retaining approximately 20 µl liquid in each well during the aspiration step to limit cell loss. Cells were washed with PBS then permeabilized for 15 minutes at room temperature with 100 µl 0.5% Triton X-100 in PBS. Cells were washed with PBS followed by Milli-Q deionized water. Cells were next treated 3 times total for 12 minutes each instance with 40 µl elution buffer which consists of 0.5M L-Glycine, 3M Urea, 3 M Guanidinium chloride, and 70 mM TCEP-HCl at pH of 2.5. Cells were washed with PBS as above. Samples were then blocked for 1 h at room temperature with 50 µl blocking buffer which consists of PBS-based Intercept buffer supplemented with 150 mM maleimide. Blocking buffer was prepared immediately prior to adding to the samples for each round. Following a PBS wash, samples were incubated overnight at 4°C with 40 µl primary antibody diluted in Intercept buffer. After overnight incubation, cells were washed with PBS then incubated for 1 hour at room temperature in 40 µl secondary antibody solution consisting of the appropriate species-specific Alexa Fluor-conjugated antibodies diluted 1:2000 in Intercept buffer. Cells were then washed with PBS and incubated with 50 µl

Hoechst 33342 diluted 1:20,000 in PBS. For the first round of imaging, cells were stained with a mixture of Hoechst and CellMask Green for 30 minutes at room temperature according to the manufacturer’s instructions. Next, cells were washed with Milli-Q water and 80 µl imaging buffer consisting of 700 mM N-Acetyl-Cysteine at pH of 7.4. Images were obtained using Operetta CLS high content imaging system (Perkin Elmer) using a 10x air objective lens. Following imaging, samples were washed with Milli-Q water after which antibodies were eluted with 3 successive 12-minute incubations at room temperature with elution buffer. Cells were washed with PBS followed by Milli-Q water. Next, 50 µl imaging buffer was added to each well and samples were imaged as above to assess removal of fluorescent signal. Cells were then washed with PBS and all steps were repeated starting at the blocking step for each round of 4i. In instances where the time between 4i rounds exceeded 3 days, following elution, the plates were fixed for 10 minutes at room temperature with 4% paraformaldehyde in PBS. In these cases, to resume staining, cells were then washed with PBS followed by Milli-Q water and treated with elution buffer lacking TCEP-HCl three times for 10 minutes, totaling 30 minutes. Afterwards, cells were washed with PBS and the next round of 4i commenced.

### Image analysis

Images were background subtracted using the rolling ball subtraction algorithm in ImageJ (2.3.0). Background-subtracted images from each round of 4i were aligned using Hoechst nuclei staining with CellProfiler (3.1.9) (McQuin et al., 2018) using the normalized cross correlation method within the Align module. Nuclei were segmented from the aligned images using the Minimum Cross Entropy thresholding method within the IdentifyPrimaryObjects module in CellProfiler. The Threshold smoothing scale and correction factor were 2.4 and 1, respectively with lower and upper threshold bounds of 0 and 1. Cell segmentation was then performed using CellMask Green staining to propagate objects from the nuclei. This was done using the Propagation method within the IdentifySecondaryObjects module. The Minimum cross entropy thresholding method was employed with a smoothing scale of 0 and correction factor of 1, the lower and upper threshold bound values set to 0 and 1, and a regularization factor of 0.05. The TrackObjects module was used to multiplex data from individual rounds of 4i. Within TrackObjects, the Follow Neighbors method was used with the maximum pixel distance of 50 and average cell diameter of 15. Comma-separated text files containing quantitative single-cell measurements of tracked objects from CellProfiler were organized using Matlab. Only objects present in every round of imaging were included in the analysis. Additional data analysis was performed using Matlab, R, and Python.

### Classifying melanoma differentiation states

To classify the differentiation state of cells based on image-based protein measurements, we generated histograms of single-cell data on each of the previously validated melanoma differentiation state markers (MITF, SOX10, NGFR and AXL) (Khaliq et al., 2021; Tsoi et al., 2018). For each protein (X), we identified an appropriate binary gate, based on which individual melanoma cells were divided into two groups of X^High^ and X^Low^ cells. The gating thresholds used on background-subtracted image data for each protein included: log(MITF) = 7.37, log(SOX10) = 6.82, log(NGFR) = 4.61, and log(AXL) = 5.60. We then used these classifications to determine the differentiation subtype of each individual melanoma cell as follows: melanocytic (M): MITF^High^ /SOX10^High^ /NGFR^Low^ /AXL^Low^; transitory (T): MITF^High^/SOX10^High^ /NGFR^High^ /AXL^Low^; neural crest-like (N): MITF^Low^ /SOX10^High^ /NGFR^High^ /AXL^High^; and undifferentiated (U): MITF^Low^ /SOX10^Low^ /NGFR^Low^ /AXL^High^; the single-cell analysis and baseline differentiation state classification were performed across 19 different melanoma cell lines representing a wide spectrum of differentiation states. To classify the differentiation state of cells in gene knockdown perturbation assays, we used a similar approach to distinguish melanocytic/transitory (MITF^High^/SOX10^High^) cells from undifferentiated (MITF^Low^ /SOX10^Low^) cells.

To classify melanoma differentiation states using bulk transcriptomic data, each melanoma cell line was assigned a series of seven differentiation signature scores, defined as the average of z-scores for the expression levels of differentiation state signature genes identified previously by Tsoi *et al* (Tsoi et al., 2018). These differentiation signatures included the four main differentiation signatures, i.e., melanocytic (M), transitory (T), neural crest-like (N) and undifferentiated (U), as well as mixtures of neighboring signatures, including melanocytic-transitory (MT), transitory-neural crest-like (TN) and neural crest-like-undifferentiated states (NU).

To determine the differentiation state of individual cells for each of the 10 melanoma cell lines profiled by single-cell RNA sequencing (Wouters et al., 2020), we used the R package AUCell (1.16.0) to quantify the enrichment of differentiation signature genes (as defined by Tsoi *et al* (Tsoi et al., 2018)) in individual cells. To minimize the impact of noise from single-cell data, we combined two or three closely related signature gene sets as follows: M-MT gene set (combination of M and MT signature genes), MT-T-TN gene set (combination of MT, T and TN signature genes), TN-N-NU gene set (combination of TN, N and NU signature genes) and NU-U set (combination of NU and U genes). We then selected cells that represented individual differentiation states based on their gated AUCell scores as follows: melanocytic cells: M-MT^High^/ TN-N-NU^Low^/NU-U^Low^; transitory cells: M-MT^Low^/ MT-T-TN^High^/ TN-N-NU^Low^/NU-U^Low^; neural crest-like cells: M-MT^Low^/ MT-T-TN^Low^/ TN-N-NU^High^/NU-U^Low^; undifferentiated cells: M-MT^Low^/ MT-T-TN^Low^/NU-U^High^. The differentiation state of individual melanoma cells derived from treatment-naïve patient tumors profiled by Tirosh *et al* (Tirosh et al., 2016) and Jerby-Arnon *et al* (Jerby-Arnon et al., 2018) were determined in the same way, except that only melanocytic and undifferentiated cells were identified and analyzed. To determine the two-class “proliferative” and “invasive” phenotypes of individual cells from the 10 cell lines and patient tumors, we used AUCell to quantify the enrichment of the two gene sets defined by Hoek *et al* (Hoek et al., 2006) for these two phenotypes at the single-cell level. We then selected cells that represent each phenotype based on their gated AUCell, including proliferative cells as proliferative^High^/ invasive^Low^ and invasive cells as proliferative^Low^/ invasive^High^.

### Random forest classification

We used random forest classification to test the predictivity of AP-1 variations for melanoma differentiation state using single-cell protein data collected by immunofluorescence imaging of 19 melanoma cell lines. We randomly sampled a total of 10,000 cells, including 2,500 from each of the four differentiation states, in a way that they represented all 19 cell lines and 4 distinctive differentiation states (melanocytic, transitory, neural crest-like and undifferentiated) as equally as possible. By random sampling, we aimed to minimize potential biases associated with genotype differences among cell lines. We used the data from 80% of the sampled cells to train a random forest classification model to predict the differentiation state of each individual cell. We then used the remaining 20% of cells to independently validate model predictions. We also evaluated the performance of random forest models in predicting differentiation states of independent cell lines using “leave-one-line-out” cross-validation. For this purpose, at each iteration, we excluded cells from one cell line, trained a model using the remaining 18 cell lines, and then used the trained model to predict the differentiation state of cells from the left-out cell line.

Model training, cross-validation and independent validation were all performed in Python (3.9.2) using the scikit-learn library (0.24.1) (Pedregosa et al., 2018). To standardize the model input, protein levels of each AP-1 measurement were normalized across cells to zero mean and unit variance (z-score scaled) using the StandardScaler() function. The random forest model was trained using the RandomForestClassifier() function. In training the full model with all 17 AP-1 factors, all parameters were as defined in the default settings, except the number of trees in the forest (n_estimators = 100) and maximum tree depth (max_depth = 14), which were separately optimized through 5-fold Stratified Shuffle Split cross-validation on the training set, using the StratifiedShuffleSplit() function with 10 times splitting iterations (n_splits = 10). In leave-one-line-out cross-validation, models built based on the top six AP-1 factors were trained using n_estimators = 500, while other parameters were the same as in the full model.

The random forest model performance was evaluated based on accuracy and Area Under the Receiver Operating Characteristic Curve (ROC AUC). Accuracy reports the fraction of correctly classified samples, i.e., true positives and true negatives, and it was calculated using the accuracy_score() function. The ROC AUC scores were calculated using the roc_auc_score() function with the One-vs-rest option (multi_class = ‘ovr’), which computes the AUC of each class against the rest. The ROC AUC scores consider both the sensitivity (true positive rate) and specificity (true negative rate) of the model predictions.

To assess the importance of each AP-1 factor in explaining the predictions made by the random forest model for each individual cell in the independent validation set, we used the SHapley Additive exPlanations (SHAP) package (Lundberg and Lee, 2017). SHAP provides a model agnostic measure of feature importance based on Shapley values, which assign importance of input features based on their contribution to the model output prediction. Mathematically, given a specific prediction output by model *f* with input *x*, Shapley value for feature *i*, φ*_i_(f,x)*, is the average of feature *i’s* marginal contributions across all possible orders of features being included (Lundberg and Lee, 2017; Lundberg et al., 2018):

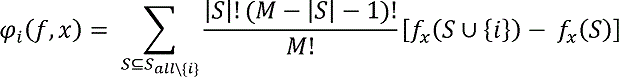

where *M* is the total number of features, *|S|* denotes number of entries in set *S* and the term is the marginal contribution of feature *i*. In SHAP, the marginal impact of a feature is defined as the change in the expected value of the model output *f(x)* when that feature is observed versus unknown:

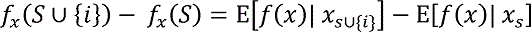

where *x_s_* is a subset of features with only set *S* is observed.

### Partial least squares regression (PLSR) modeling

We used PLSR analysis to test whether the relationships between the patterns of AP-1 gene expression and melanoma differentiation state were recapitulated at the transcriptional level. Bulk RNA sequencing data of 53 melanoma cell lines used for PLSR analysis were obtained from Tsoi *et al* (Tsoi et al., 2018). We first log_2_-transformed the gene expression data (reported as FPKM) with an offset of 1. Input vectors for PLSR analysis were then created by combining the z-scored expression data for fifteen AP-1 transcription factor family genes, including FOS, FOSL1, FOSL2, FOSB, JUN, JUNB, JUND, ATF1, ATF2, ATF3, ATF4, ATF5, ATF6, ATF6B and ATF7, across 53 cell lines. The response variables for each cell line were then assembled as a series of seven signature scores, defined as the average of z-scores for the expression levels of differentiation state signature genes (Tsoi et al., 2018). The PLSR model was trained in python using the scikit-learn library and the PLSRegression() function. To evaluate the predictability of the linear relationship between the input and output variables using the same dataset, we used leave-one-out cross-validation by LeaveOneOut() function. To independently validate the model, we used RNA sequencing data from an independent panel of 32 melanoma cell lines in the Cancer Cell Line Encyclopedia (CCLE) (Ghandi et al., 2019). As with the training dataset, we first log_2_-transformed the CCLE gene expression data (reported as RPKM) with an offset of 1. We then created input vectors by combining the z-scored expression data for fifteen AP-1 transcription factor family genes and used them in the optimized PLSR model (using the first four PLS components) trained against the original set of 53 cell lines to predict the differentiation signature scores in the new set of 32 cell lines.

The PLSR model performance was evaluated in terms of fraction of variance explained (R^2^) or predicted (Q^2^) using the explained_variance_score() function. We assessed the relative importance of each AP-1 factor in the PLSR model based on the variable importance in projection (VIP) scores, computed for the first four PLS components, at which the PLSR model achieves its optimal performance (Wold, 1994). To help interpret the directionality of the contribution, we multiplied the VIP score for each AP-1 factor by the sign of Pearson correlation coefficient between its expression levels and differentiation signature z-scores.

We compared the performance (based on 10-fold cross validation) of optimized PLSR model, built based on the top eight AP-1 genes (FOS, FOSL1, FOSL2, JUN, JUNB, JUND, ATF2 and ATF4) with optimized models built using combinations of eight randomly chosen basic leucine zipper (bZIP) transcription factors (Vinson et al., 2002) or eight randomly chosen transcription factors (Van de Sande et al., 2020) (excluding those that were explicitly involved in the differentiation signature genes) by computing empirical *P* values using 100,000 and 500,000 iterations, respectively.

To identify dynamic patterns of AP-1 changes that are associated with drug-induced changes in differentiation state, we constructed a PLSR model to relate DMSO-normalized changes in AP-1 proteins to DMSO-normalized changes in differentiation state. The input vector consists of DMSO-normalized population measurements of the seventeen AP-1 proteins, including cFOS, p-cFOS, FRA1, p-FRA1, FRA2, cJUN, p-cJUN, JUNB, JUND, p-ATF1, ATF2, p-ATF2, ATF3, ATF4, p-ATF4, ATF5 and ATF6, at 24 h and 72 h, z-scored across two MAPK inhibitor treatment conditions and eighteen cell lines. DMSO-normalized AP-1 measurements were calculated as the log ratios of drug-treated AP-1 levels relative to the DMSO control. The response vector is composed of the DMSO-normalized changes in differentiation state, which is calculated as 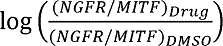. Model training, cross-validation and performance evaluation were performed the same way as in the gene-expression PLSR model. VIP scores were computed for the first three PLS components.

### Uniform manifold approximation and projection (UMAP)

UMAP was performed in R using the umap package (0.2.7.0). For single-cell protein data, we first performed principal component analysis (PCA) using the prcomp() function on the z-scored log-transformed data and selected the PCA scores from the first four principal component for UMAP analysis. The parameters used in generating the UMAP for single-cell protein data include nearest neighbor (n_neighbors) = 90, minimum distance (min_dist) = 0.7 and distance metric (metric) = Euclidean.

### Hierarchical clustering

Unsupervised hierarchical clustering of population-averaged AP-1 protein measurements was carried out in R using the stats (4.1.2) package. Clustering was performed using the hclust() function with the average algorithm as the agglomeration method. The distance matrix used for clustering was evaluated using the dist() function, with Pearson’s correlation as the distance metric.

### Single-cell regulatory network inference and clustering (SCENIC)

For the SCENIC analysis of melanoma cell lines, the baseline regulon activities inferred by the SCENIC workflow (Aibar et al., 2017; Van de Sande et al., 2020) were obtained from the .loom file published by Wouters *et al* (Wouters et al., 2020). The .loom file was imported to R for downstream analysis using the SCopeLoomR package (0.13.0).

Single-cell RNA sequencing data for patient-derived melanoma tumors were obtained from previous studies published by Tirosh *et al* (Tirosh et al., 2016) and Jerby-Arnon *et al* (Jerby-Arnon et al., 2018). Single-cell gene expression analysis and SCENIC was focused on 2072 malignant melanoma cells, which were distinguished (by the authors) from non-malignant cells based on gene copy number variations. For quality control, we first selected 14,689 genes which were detected in more than 1% of the cells (i.e., 20 cells) with at least 103 logged TPM counts, using the single-cell analysis toolkit Scanpy (1.7.1) (Wolf et al., 2018) in Python. Using this dataset, we then inferred regulons using pySCENIC (0.11.0) in a Nextflow pipeline adapted from Wouters *et al* (Wouters et al., 2020), performing 100 SCENIC runs on the data. As in Wouters *et al*’s regulon filtering criteria, only regulons that had more than 10 target genes and recurred in at least 80/100 runs were retained. Target genes (used in AUCell calculation) that appear in at least 80% of the runs in regulons that recurred 100 times, and all target genes for regulons that recurred 80-100 times were retained. This analysis pipeline resulted in 373 motif regulons.

### Partial correlation analysis

Partial correlation analysis is used for the evaluation of correlations between pairs of variables while controlling for the variance explained by a third variable. We used pairwise partial correlation analysis to evaluate correlations between changes induced by MAPK inhibitors in each of the AP-1 protein levels and p-ERK levels across different cell lines, while controlling for the baseline (drug-naïve) variance of AP-1 levels across the same cell lines. AP-1 and p-ERK data were averaged across two replicates and log-transformed. To assess if any of the AP-1 factors would capture drug-specific changes in ERK signaling, we then used the Matlab function partialcorr() to evaluate the Pearson’s partial correlation coefficients (and the associated *P* values) between p-ERK and AP-1 levels across cell lines for each MAPK inhibitor treatment condition, while correcting for differences in their baseline (DMSO condition) AP-1 levels.

## QUANTIFICATION AND STATISTICAL ANALYSIS

Single-cell protein abundance was quantified from microscopy images using CellProfiler (3.1.9). No statistical method was used to predetermine sample size. Sample sizes were chosen based on similar studies in the relevant literature. The experiments were not randomized. The investigators were not blinded to allocation during experiments and outcome assessment. All boxplots and violin plots highlight the median, lower and upper quartiles. Whiskers in boxplots indicate 1.5 times interquartile ranges. Sample size (i.e., number of cells or replicates) are indicated in the figure legend. The significance of pairwise correlations were evaluated based on *P* values associated with the corresponding two-sided Pearson’s correlation analysis. Statistical significance of changes in population-averaged protein abundance across different drugs and/or timepoints were determined based on one-way or two-way analysis of variance (ANOVA), as indicated in the figure legends. To identify the statistical significance of differences between mean of measurements of two different groups, *P* values were determined using paired or unpaired two-sided *t* test, as indicated in the figure legends. Statistical analyses were performed using MATLAB (2020b) and R (4.0.4).

### Supplemental Datasets

**Dataset S1.** Baseline (drug-naïve) single-cell protein levels of 17 AP-1 proteins and four melanoma differentiation state markers MITF, SOX10, NGFR and AXL, measured by 4i across 19 BRAF-mutant melanoma cell lines. Protein data shown for each condition and individual cell represents the log-transformed values measured in the presence of DMSO (vehicle) at 24 h.

**Dataset S2.** Single-cell enrichment of Tsoi *et al* differentiation signature gene sets (measured by AUCell): M-MT gene set (combination of M and MT signature genes), MT-T-TN gene set (combination of MT, T and TN signature genes), TN-N-NU gene set (combination of TN, N and NU signature genes) and NU-U set (combination of NU and U genes), as well as Hoek *et al* signature gene sets: proliferative and invasive, on 10 melanoma cell lines profiled by Wouters *et al*.

**Dataset S3**. Single-cell AP-1 regulon activities, enrichment of M-MT gene set, MT-T-TN gene set, TN-N-NU gene set, NU-U gene set, proliferative gene set, and invasive gene set on 11 treatment-naïve patient tumor samples profiled by Tirosh *et al* and Jerby-Arnon *et al*.

**Dataset S4.** Single-cell protein levels of 17 AP-1 proteins, differentiation state markers MITF and NGFR, and p-ERK levels measured by 4i across 18 BRAF-mutant melanoma cell lines following exposure to either vehicle (DMSO), BRAF inhibitor (vemurafenib at 0.316 µM), or the combination of BRAF and MEK inhibitors (vemurafenib at 0.316 µM and trametinib at 0.0316 µM) for 24 and 72 h. Protein data shown for each condition and individual cell represents the log-transformed values.

**Dataset S5.** Population-averaged protein levels of 17 AP-1 proteins, differentiation state markers MITF and NGFR, and p-ERK levels measured by 4i across 18 BRAF-mutant melanoma cell lines following exposure to either vehicle (DMSO), BRAF inhibitor (vemurafenib at 0.316 µM), or the combination of BRAF and MEK inhibitors (vemurafenib at 0.316 µM and trametinib at 0.0316 µM) for 24 and 72 h (in 3 replicates). Protein data shown for each condition represents the log-transformed values.

**Dataset S6.** Population-averaged protein levels of five AP-1 proteins and differentiation state markers MITF and SOX10 measured by 4i across siRNA knockdown conditions in COLO858, C32 and LOXIMVI at 96 h. Protein data shown for each condition represents the log-transformed values.

