## Supplemental Information for "AP-1 transcription factor network explains diverse patterns of cellular plasticity in melanoma cells"

**Figure S1. Single-cell AP-1 protein levels predict differentiation state heterogeneity in melanoma cells. Related to Figure 1.**

**(A)** Single-cell distributions of seventeen AP-1 factors and four differentiation state markers measured across 19 cell lines and shown by violin plots highlighting the median and interquartile (25% and 75%) ranges. **(B)** Random Forest model cross-validation performance (using the top six AP-1 factors) to predict the differentiation state of new cells from independent cell lines not included in model training. At each iteration, one cell line was removed, a model was built using randomly sampled cells from the remaining 18 cell lines, and then the trained model was used to predict the differentiation state of randomly selected cells from the left-out cell line. Red dash line indicates accuracy from random prediction (25%). **(C)** Confusion matrix showing the cross-validation performance of the random forest classifier. Numbers shown are in terms of average percentage of cells in the indicated categories. **(D)** Confusion matrices showing the model cross-validation performance for each independent cell line.

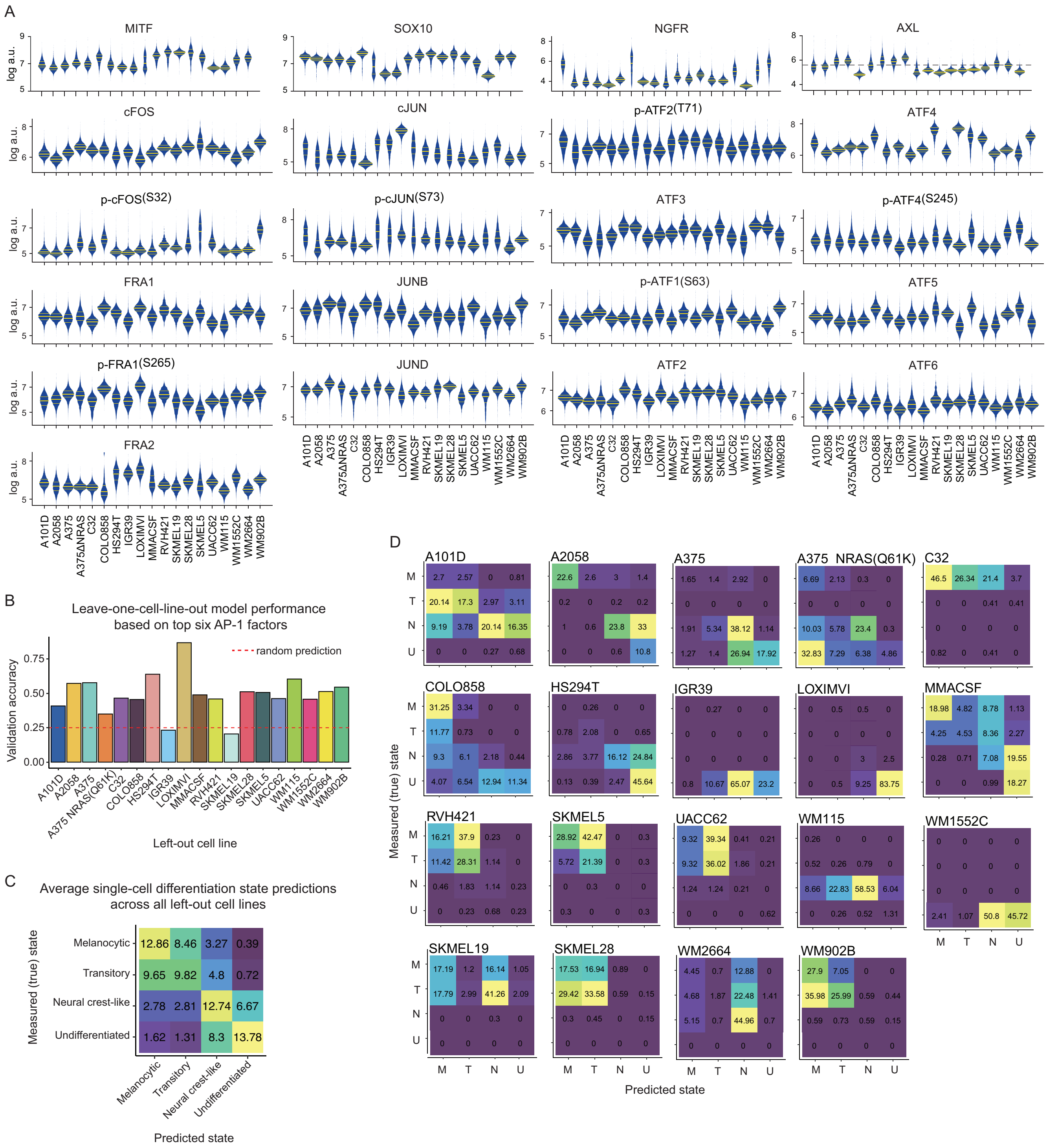
A
MITF
SOX10
NGFR
AXL
cFOS
cJUN
p-ATF2(T71)
ATF4
p-cFOS(S32)
p-cJUN(S73)
ATF3
p-ATF4(S245)
FRA1
JUNB
p-ATF1(S63)
ATF5
p-FRA1(S265)
JUND
ATF2
ATF6
FRA2
log a.u.
A101D A2058 A375 A375ΔNRAS C32 COLO858 HS294T IGR39 LOXIMVI MMACSF RVH421 SKMEL19 SKMEL28 SKMEL5 UACC62 WM115 WM1552C WM2664 WM902B
B
Leave-one-cell-line-out model performance based on top six AP-1 factors
Validation accuracy
random prediction
Left-out cell line
C
Average single-cell differentiation state predictions across all left-out cell lines
Measured (true) state
Melanocytic
Transitory
Neural crest-like
Undifferentiated
Predicted state
D
Measured (true) state
Predicted state
M T N U

**Figure S2. AP-1 protein, transcript levels and activities predict variations in differentiation state and phenotype across melanoma cell lines and single cells. Related to Figures 1-3. (A**) Summary of contributions of 15 AP-1 factors in predicting each melanoma differentiation state based on analyses performed in this study on single-cell protein measurements and bulk gene expression data. Significant positive and negative predictors are highlighted. Significant AP-1 proteins were determined based on their SHAP importance in the random forest model. Significant AP-1 genes were determined based on the significance of their VIP scores (i.e., magnitude greater than 1) in the PLSR model. **(B-F)** Single-cell distributions of the activity of SCENIC regulons for FOSL2 **(B)**, JUN **(C)**, FOSL1 **(D)**, and FOS **(E)** motifs, as well as the ratio of FOS and JUN regulon activities **(F),** measured using AUCell in individual cells (from 10 melanoma cell lines profiled by Wouters *et al*) across the two-class ("proliferative" versus "invasive") melanoma phenotypes. The phenotype of individual cells was determined based on their gated levels of enrichment (quantified by AUCell) for the gene signatures as defined by Hoek *et al*. **(G-K)** Single-cell distributions of the AUCell activity of SCENIC regulons for FOSL2 **(G)**, JUN **(H)**, FOSL1 **(I)**, and FOS **(J)** motifs, as well as the ratio of FOS and JUN regulon activities **(K)**, quantified in individual cells from 12 treatment-naïve melanoma tumors as profiled by Tirosh *et al* and Jerby-Arnon *et al*. Statistical comparisons were performed using two-sided unpaired *t* test. Boxplot hinges correspond to the lower and upper quartiles, with a band at median. Whiskers indicate 1.5 times interquartile ranges.

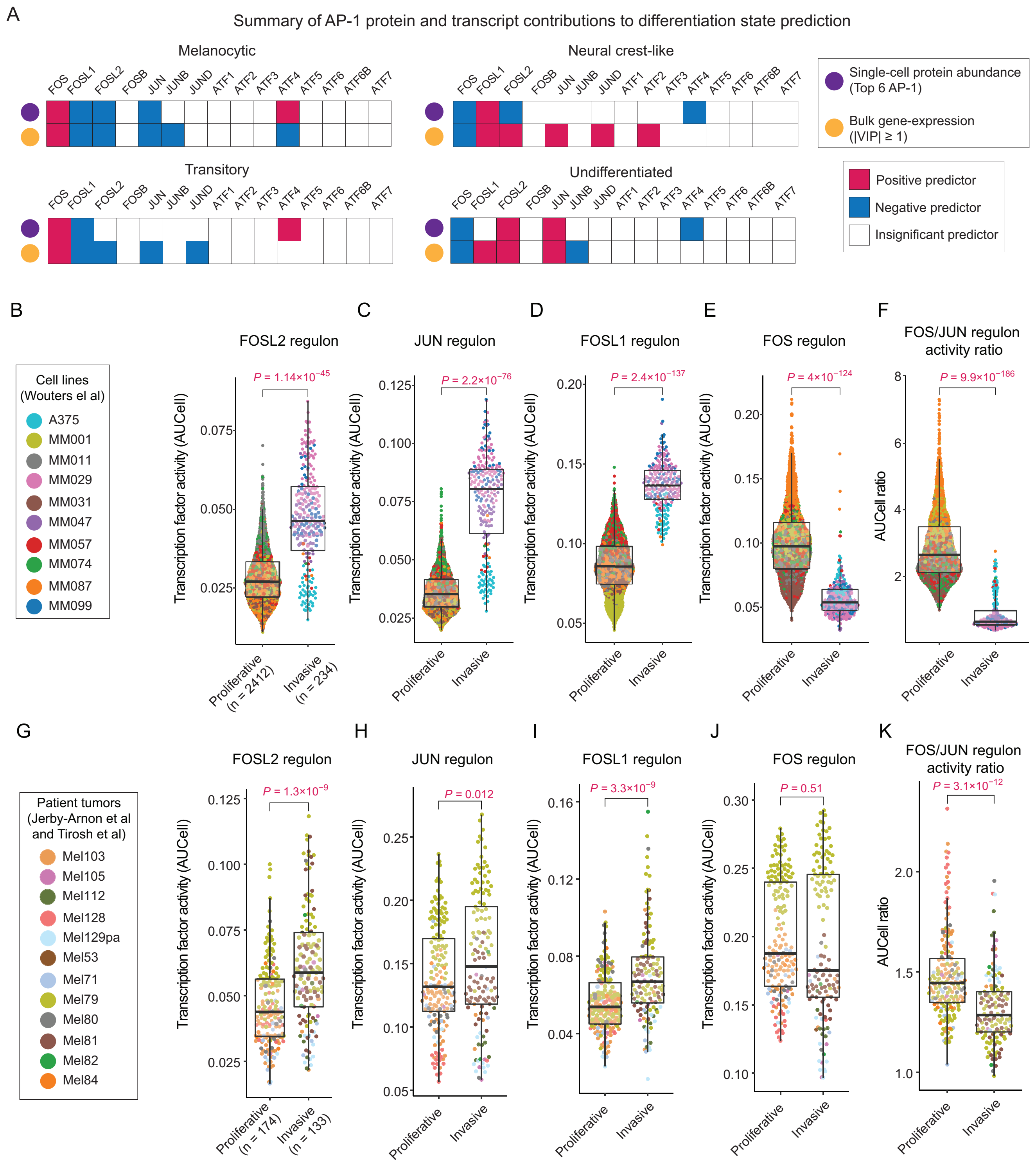

A
Summary of AP-1 protein and transcript contributions to differentiation state prediction
Melanocytic
Neural crest-like
Transitory
Undifferentiated
FOS FOSL1 FOSL2 FOSB JUN JUNB JUND ATF1 ATF2 ATF3 ATF4 ATF5 ATF6 ATF6B ATF7
Single-cell protein abundance (Top 6 AP-1)
Bulk gene-expression (|VIP| ≥ 1)
Positive predictor
Negative predictor
Insignificant predictor
B
C
D
E
F
Cell lines (Wouters el al)
A375
MM001
MM011
MM029
MM031
MM047
MM057
MM074
MM087
MM099
FOSL2 regulon
JUN regulon
FOSL1 regulon
FOS regulon
FOS/JUN regulon activity ratio
P = 1.14×10−45
P = 2.2×10−76
P = 2.4×10−137
P = 4×10−124
P = 9.9×10−186
Transcription factor activity (AUCell)
AUCell ratio
Proliferative (n = 2412)
Invasive (n = 234)
Proliferative
Invasive
G
H
I
J
K
Patient tumors (Jerby-Arnon et al and Tirosh et al)
Mel103
Mel105
Mel112
Mel128
Mel129pa
Mel53
Mel71
Mel79
Mel80
Mel81
Mel82
Mel84
P = 1.3×10−9
P = 0.012
P = 3.3×10−9
P = 0.51
P = 3.1×10−12
Proliferative (n = 174)
Invasive (n = 133)

**Figure S3. AP-1 protein measurements (including FOS and JUN subfamily members), measured in 18 cell lines before and following treatment with MAPK inhibitors for 24 and 72 h. Related to Figures 4 and 5.** Bar height indicates mean values between two replicates shown as black dots. *P*-values show the statistical significance of treatment condition effect on indicated protein levels based on one-way ANOVA test.

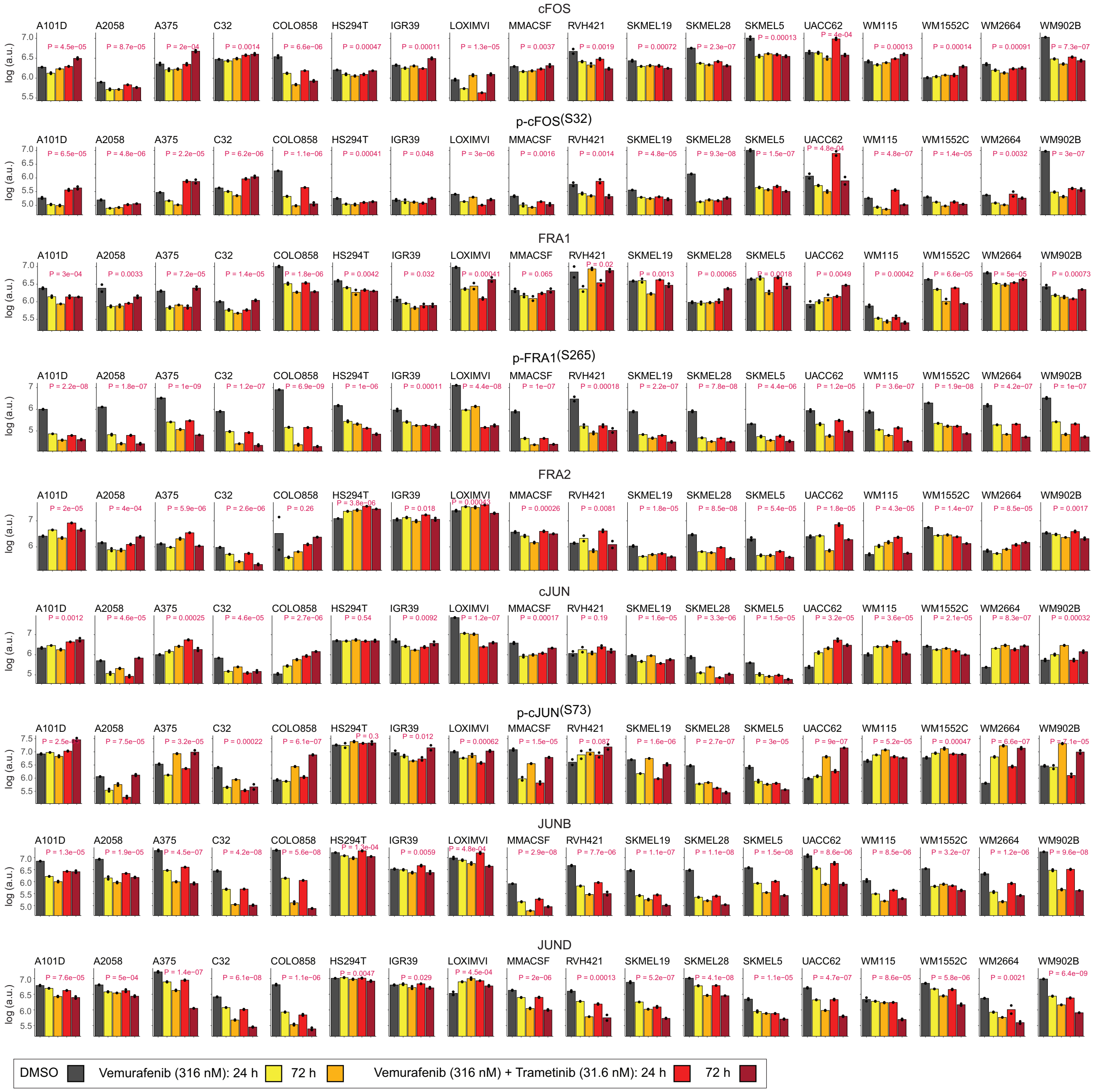

cFOS
p-cFOS(S32)
FRA1
p-FRA1(S265)
FRA2
cJUN
p-cJUN(S73)
JUNB
JUND
A101D
A2058
A375
C32
COLO858
HS294T
IGR39
LOXIMVI
MMACSF
RVH421
SKMEL19
SKMEL28
SKMEL5
UACC62
WM115
WM1552C
WM2664
WM902B
log (a.u.)
DMSO
Vemurafenib (316 nM): 24 h
72 h
Vemurafenib (316 nM) + Trametinib (31.6 nM): 24 h
72 h

**Figure S4. AP-1 protein measurements (including ATF subfamily members) and differentiation state markers MITF and NGFR, measured in 18 cell lines before and following treatment with MAPK inhibitors for 24 and 72 h. Related to Figures 4 and 5.** Bar height indicates mean values between two replicates shown as black dots. *P*-values show the statistical significance of treatment condition effect on indicated protein levels based on one-way ANOVA test.

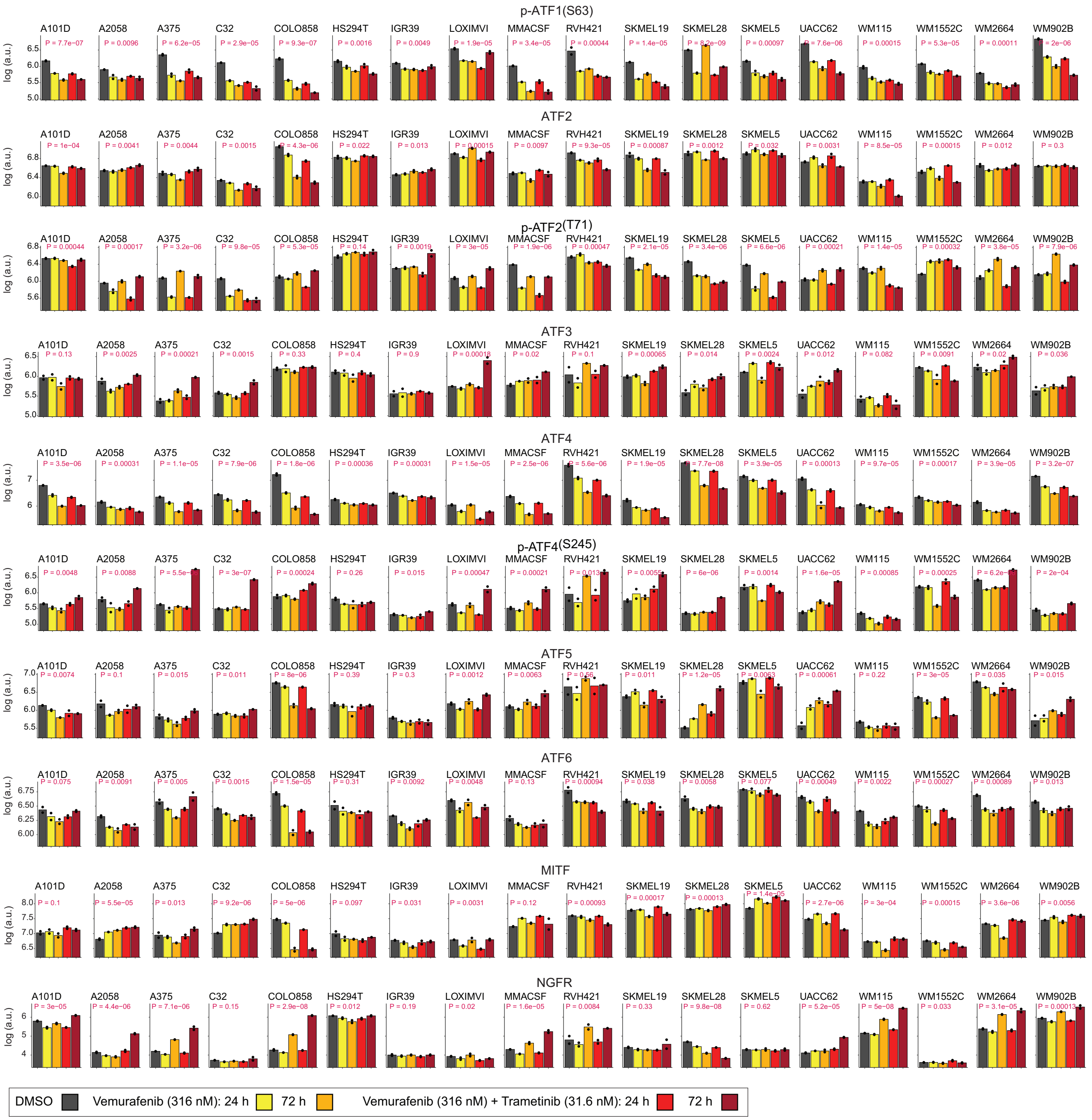

p-ATF1(S63)
ATF2
p-ATF2(T71)
ATF3
ATF4
p-ATF4(S245)
ATF5
ATF6
MITF
NGFR
log (a.u.)
A101D
A2058
A375
C32
COLO858
HS294T
IGR39
LOXIMVI
MMACSF
RVH421
SKMEL19
SKMEL28
SKMEL5
UACC62
WM115
WM1552C
WM2664
WM902B
DMSO
Vemurafenib (316 nM): 24 h
72 h
Vemurafenib (316 nM) + Trametinib (31.6 nM): 24 h
72 h

**Figure S5. Representative immunofluorescence images of AP-1 proteins, differentiation state markers and p-ERK levels following MAPK inhibitor treatment. Related to Figures 5, 6. (A)** Immunofluorescence images of AP-1 proteins cFOS, p-cFOS, cJUN, p-cJUN, FRA1, p-FRA1 and differentiation state markers MITF and NGFR in COLO858 cells before and after treatment with indicated inhibitors at indicated doses and timepoints. Each experiment was repeated twice with similar result. Scale bars represent 100 µm. **(B)** Immunofluorescence images of p-ERK in representative cell lines (LOXIMVI, COLO858 and SKMEL28) before and after treatment with indicated inhibitors at indicated doses and timepoints. Each experiment was repeated twice with similar result. Scale bars represent 100 µm.

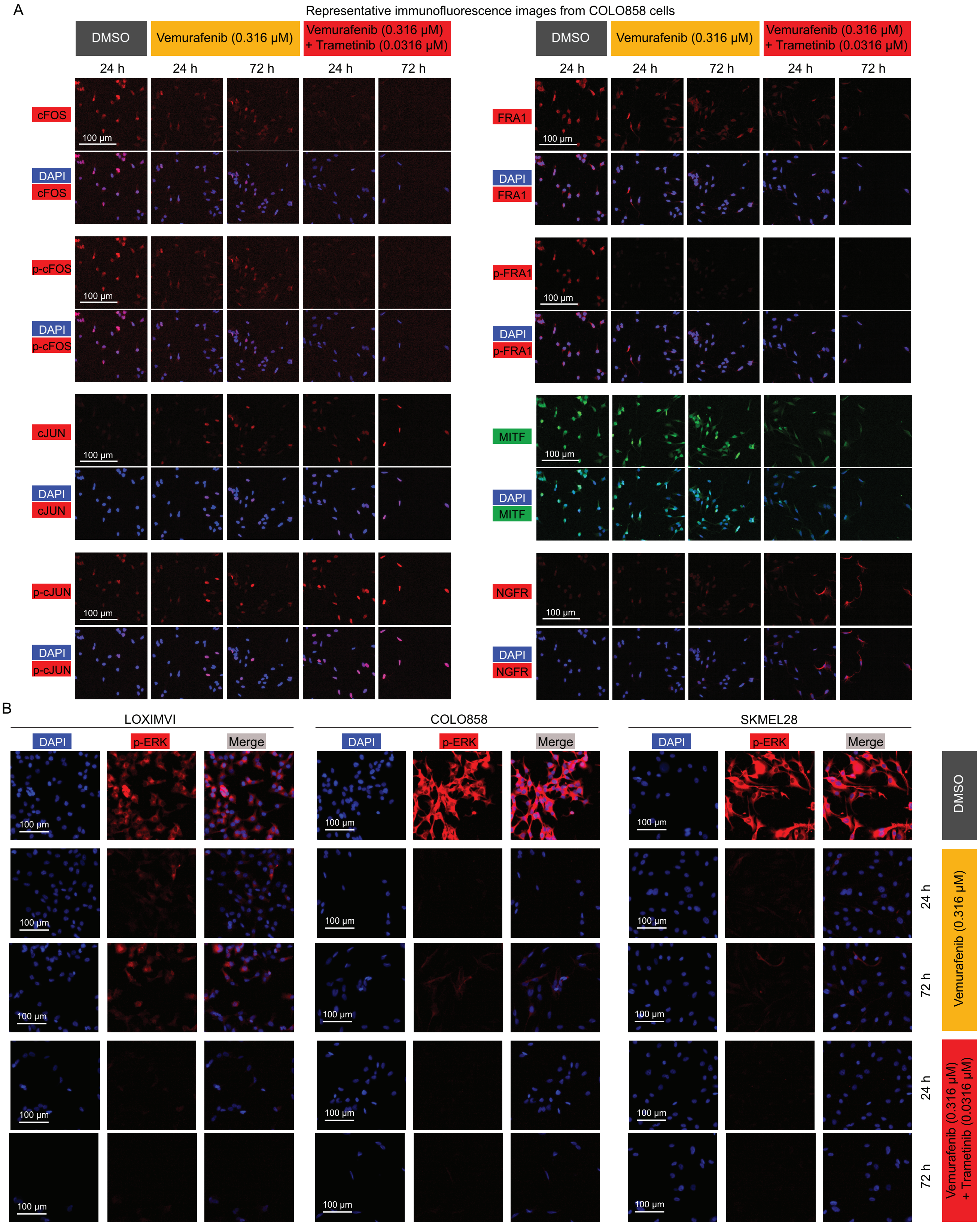

A
Representative immunofluorescence images from COLO858 cells
DMSO
Vemurafenib (0.316 µM)
Vemurafenib (0.316 µM) + Trametinib (0.0316 µM)
24 h
72 h
cFOS
DAPI
p-cFOS
cJUN
p-cJUN
FRA1
p-FRA1
MITF
NGFR
100 µm
B
LOXIMVI
COLO858
SKMEL28
DAPI
p-ERK
Merge

**Figure S6. Correlation analysis between drug-naïve (baseline) p-ERK and AP-1 protein levels across melanoma cell lines. Related to Figure 6. (A-B)** Pearson's correlations (evaluated across 19 drug-naïve cell lines) and associated *P* values between each of the 17 AP-1 measurements and p-ERK levels. Each data-point in **(B)** represents population-averaged measurements across all cells pooled from two replicates for each cell line.

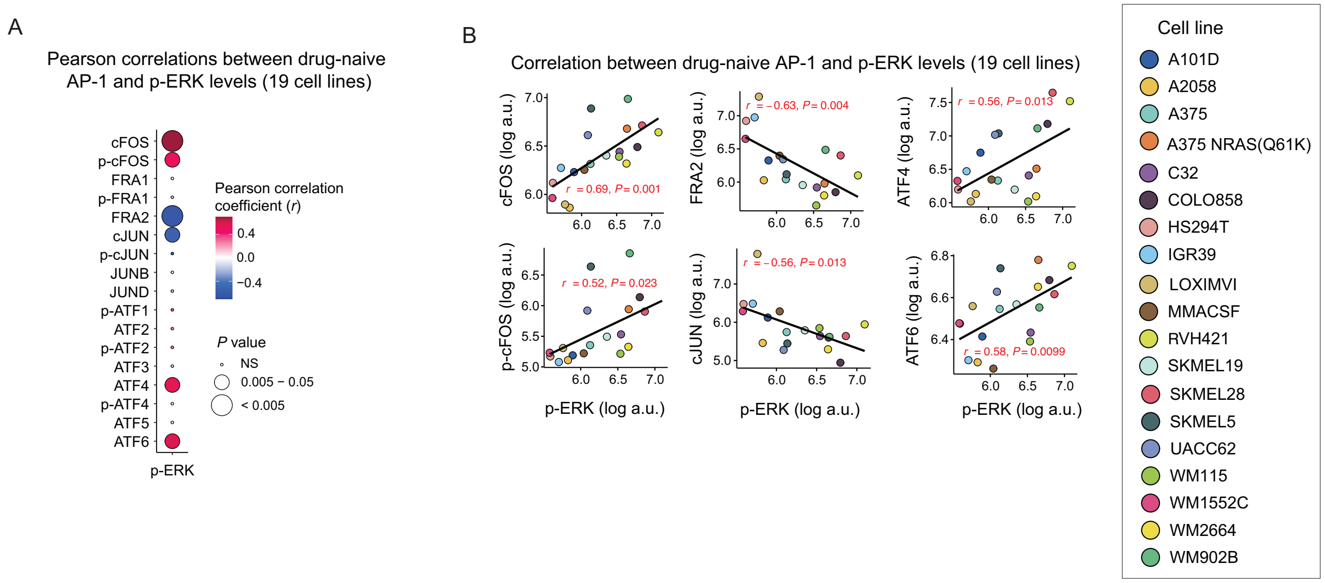

A
Pearson correlations between drug-naive AP-1 and p-ERK levels (19 cell lines)
cFOS
p-cFOS
FRA1
p-FRA1
FRA2
cJUN
p-cJUN
JUNB
JUND
p-ATF1
ATF2
p-ATF2
ATF3
ATF4
p-ATF4
ATF5
ATF6
p-ERK
Pearson correlation coefficient (r)
0.4
0.0
−0.4
P value
NS
0.005 − 0.05
< 0.005
B
Correlation between drug-naive AP-1 and p-ERK levels (19 cell lines)
cFOS (log a.u.)
r = 0.69, P = 0.001
FRA2 (log a.u.)
r = −0.63, P = 0.004
ATF4 (log a.u.)
r = 0.56, P = 0.013
p-cFOS (log a.u.)
r = 0.52, P = 0.023
cJUN (log a.u.)
r = −0.56, P = 0.013
ATF6 (log a.u.)
r = 0.58, P = 0.0099
p-ERK (log a.u.)
Cell line
A101D
A2058
A375
A375 NRAS(Q61K)
C32
COLO858
HS294T
IGR39
LOXIMVI
MMACSF
RVH421
SKMEL19
SKMEL28
SKMEL5
UACC62
WM115
WM1552C
WM2664
WM902B

**Figure S7. Representative immunofluorescence images of AP-1 proteins, differentiation state markers following different AP-1 knockdown conditions. Related to Figures 7.**

Immunofluorescence images (by 4i) of AP-1 proteins cFOS, FRA2, cJUN, JUND, and differentiation state markers MITF and SOX10 in COLO858 cells before and after treatment with indicated siRNA conditions for 96 h. Each experiment was repeated three times with similar result. Scale bars represent 100 μm.

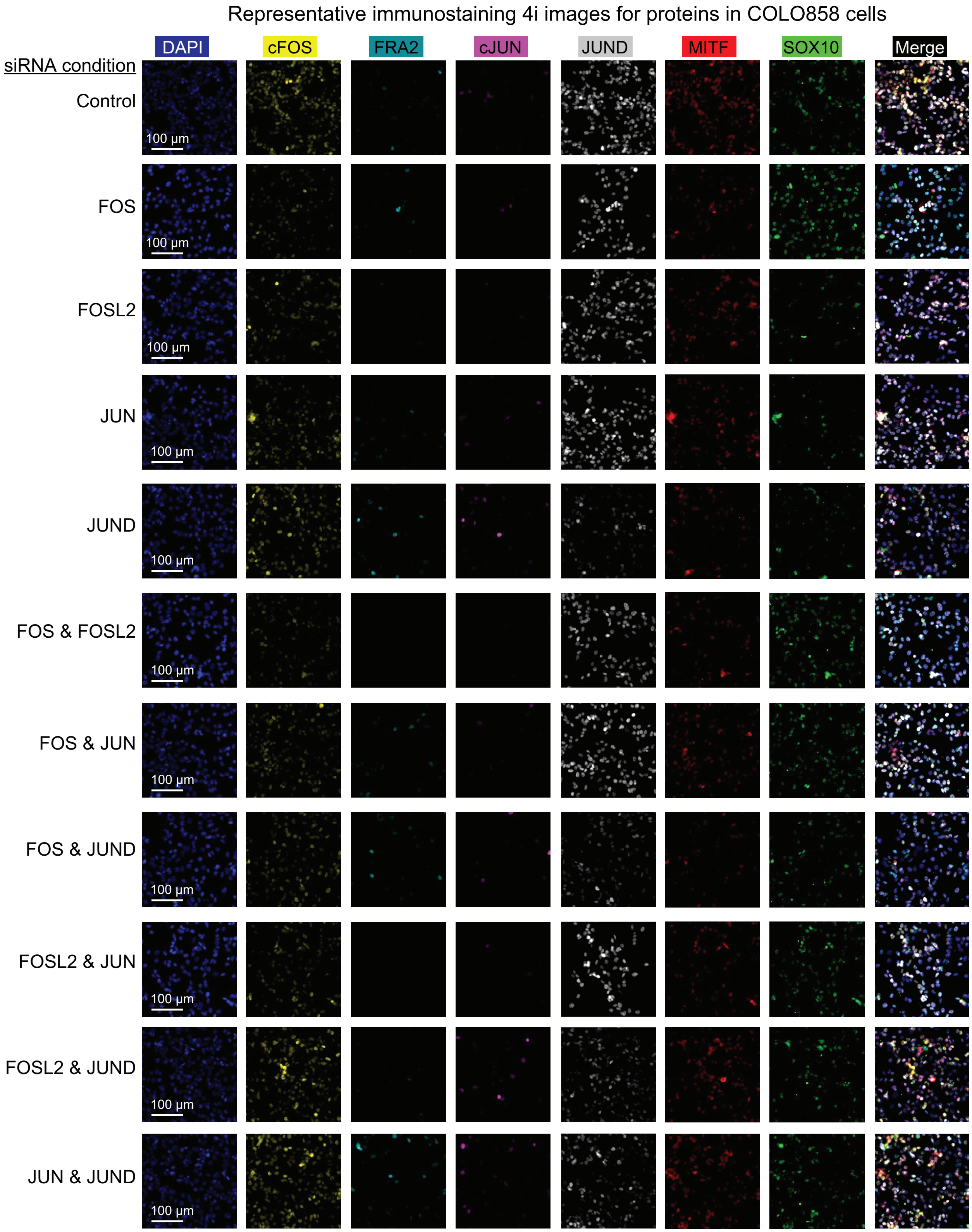
Representative immunostaining 4i images for proteins in COLO858 cells
DAPI
cFOS
FRA2
cJUN
JUND
MITF
SOX10
Merge
siRNA condition
Control
FOS
FOSL2
JUN
JUND
FOS & FOSL2
FOS & JUN
FOS & JUND
FOSL2 & JUN
FOSL2 & JUND
JUN & JUND
100 μm

**Table S1.** The siRNA sequences used in this study.

| Target Gene | Dharmacon Cat# | siRNA sequence |
|---|---|---|
| **Non-targeting control** | D-001810-10-05 | UGGUUUACAUGUCGACUAA,<br>UGGUUUACAUGUUGUGUGA,<br>UGGUUUACAUGUUUUCUGA,<br>UGGUUUACAUGUUUUCCUA |
| **FOS** | J-003265-09 | GGGAUAGCCUCUCUUACUA |
| | J-003265-10 | ACAGUUAUCUCCAGAAGAA |
| | J-003265-12 | GCAAUGAGCCUUCCUCUGA |
| **FOSL1** | J-004341-06 | GAGCUGCAGUGGAUGGUAC |
| | J-004341-07 | AAUCUGGGCUGCAGCGAGA |
| | J-004341-08 | GAGUAAGGCGCGAGCGGAA |
| **FOSL2** | J-004110-13 | GGCCCAGUGUGCAAGAUUA |
| **cJUN** | J-003268-10 | GAGCGGACCUUAUGGCUAC |
| | J-003268-12 | GAAACGACCUUCUAUGACG |
| **JUND** | J-003900-12 | GAAACACCCUUCUACGGCG |
| | J-003900-13 | CCGACGAGCUCACAGUUCC |
| | J-003900-14 | UCAAGAGUCAGAACACGGA |
| | J-003900-15 | GUUCGAUUCUGCCCUAUUU |
